# Cited4a limits cardiomyocyte dedifferentiation and proliferation during zebrafish heart regeneration

**DOI:** 10.1101/2024.12.05.626917

**Authors:** Rachel Forman-Rubinsky, Wei Feng, Brent T. Schlegel, Angela Paul, Daniel Zuppo, Katarzyna Kedziora, Donna Stoltz, Simon Watkins, Dhivyaa Rajasundaram, Guang Li, Michael Tsang

**Affiliations:** Department of Cell Biology, University of Pittsburgh, School of Medicine, Pittsburgh, PA; Center for Integrative Organ Systems, University of Pittsburgh, School of Medicine, Pittsburgh, PA; Department of Pediatrics, Division of Health Informatics, Children’s Hospital of Pittsburgh, Pittsburgh, PA; Center for Biological Imaging, University of Pittsburgh, School of Medicine, Pittsburgh, PA

**Author notes:** Department of Medicine, University of Rochester Medical Center Rochester, NY.

## Abstract

Cardiac regeneration involves the interplay of complex interactions between many different cell types, including cardiomyocytes. The exact mechanism that enables cardiomyocytes to undergo dedifferentiation and proliferation to replace lost cells has been intensely studied. Here we report a single nuclear RNA sequencing profile of the injured zebrafish heart and identify distinct cardiomyocyte populations in the injured heart. These cardiomyocyte populations have diverse functions, including stress response, myofibril assembly, proliferation and contraction. The contracting cardiomyocyte population also involves the activation of maturation pathways as an early response to injury. This intriguing finding suggests that constant maintenance of a distinctive terminally differentiated cardiomyocyte population is important for cardiac function during regeneration. To test this hypothesis, we determined that *cited4a,* a p300/CBP transcriptional coactivator, is induced after injury in the mature cardiomyocyte population. Moreover, loss-of-*cited4a* mutants presented increased dedifferentiation, proliferation and accelerated heart regeneration. Thus, suppressing cardiomyocyte maturation pathway activity in injured hearts could be an approach to promote heart regeneration.

## Introduction

Myocardial infarction (MI) leaves the heart permanently damaged and more susceptible to future cardiac failure [1–3]. One reason is that mammalian adult cardiomyocytes (CMs) are in a post-mitotic, non-proliferative state and therefore fail to replace the lost CMs [2, 3]. During embryonic and early postnatal development, mammalian CMs are in a mononucleated, diploid state and proliferate extensively before completing a final round of division and exiting the cell cycle [4, 5]. During this transition in mice, 95% of CMs stop dividing and become a binucleated or mononucleated polyploid cells [5–7]. In contrast, adult zebrafish CMs remain mononucleated, diploid cells with the ability to proliferate and can fully regenerate following injury, including revascularization and the resolution of fibrotic scar tissue [8–10]. The zebrafish offers an opportunity to investigate the molecular mechanisms that regulate heart regeneration.

Although adult mammals lack the ability to regenerate myocardial tissue after injury, robust heart regeneration has been reported in neonatal mice [11]. In zebrafish and neonatal mice, new CMs are formed from existing CMs through dedifferentiation, proliferation, and redifferentiation/maturation [12–14]. Prior to proliferation, zebrafish CMs at the injury border zone dedifferentiate into an immature transitional state exhibiting structural, metabolic, and electrophysiological changes [13–20]. The ability of CMs to dedifferentiate in response to injury is key to their regenerative ability. Genetic fate-mapping and single-cell RNA sequencing from neonatal mice and zebrafish have shown that only a small percentage of CMs near the injury border zone undergo this process after injury [15, 16, 21]. How specific CMs maintain their mature state while others are activated to enter the cell cycle and then return to mature CM function following regeneration in the spatial context of the injured myocardium is not well established.

To identify molecular events that induce or restrict proliferation, we generated a single-nucleus RNA expression atlas of the zebrafish ventricle during early heart regeneration. Our findings confirm the cellular heterogeneity of CMs in regeneration shown by other studies in mice and zebrafish [14, 15, 21–24]. Eight distinct subpopulations of CMs, including injured, dedifferentiating, proliferating, and mature cells, were identified at 3 days post amputation (dpa) that include injured, dedifferentiating, proliferating, and mature cells. The finding that CM maturation pathways are active at 3 dpa is intriguing as it suggests that in response to injury, distinct CM populations maintain their differentiated state via the activation of maturation genes. We identified *cited4a* (CREB-binding protein (CBP)/p300 interacting transactivator with ED (glutamic acid/aspartic acid)-rich tail 4a), as a gene with increased expression in mature CMs following injury. We speculated that *cited4a* may function in balancing CM maturation and proliferation during heart regeneration. *cited4a* mutant zebrafish presented increased dedifferentiation marker gene expression and increased CM proliferation, which was accompanied with accelerated fibrotic tissue resolution after ventricular amputation.

Our findings suggest that the activation of CM maturation is required during the early phases of regeneration to prevent a population of CMs from entering the cell cycle. This type of injury response would be important for the damaged heart to maintain its primary ability to contract and circulate blood while recovering from cardiac injury. Therefore, suppressing the CM maturation pathway early could increase CM dedifferentiation and proliferation during regeneration. Understanding the dynamics of CM populations in injury will reveal new approaches to stimulate regeneration in the human heart.

## RESULTS

### snRNA-seq reveals distinct CM clusters

scRNA-seq studies on adult zebrafish heart regeneration have been reported [23]. However, CMs are underrepresented because their cell size hinders their capture and therefore the exact CM population during regeneration has not been clearly described. To profile adult zebrafish heart regeneration at a single cell resolution, we therefore performed snRNA-seq on uninjured adult hearts and at 3 dpa (**Fig. 1A**). Using Seurat, unsupervised graph-based clustering was performed to pool the 20151 nuclei from uninjured (8245) and 3 dpa (11906) samples to allow for comparable cluster identification across these two integrated samples (**Supplementary Fig. 1A**). The cell clusters were visualized through uniform manifold approximation and projection (UMAP) and a total of 15 distinct cell clusters (0-14) were identified (**Fig. 1B**, **Supplementary Fig. 1B,** and **Supplementary Table S1**). Differential gene expression of each cluster relative to all other clusters revealed the top markers that define the 15 clusters and revealed endothelial cells (*spock3*), megakaryocytes (*itgb3b*), macrophages (*marco*) and lymphoid cells (*cd74*) (**Fig. 1 C, D**, and **Supplementary Fig. 1C, D**). In addition, endocardial cells (*bcam*), erythrocytes *(hbba1*), epicardial cells (*tbx18)*, and pericytes (*pdgfrb)* were identified (**Supplementary Fig. 1C, D**). These marker genes for these populations have been previously used to define these cell types in regenerating adult zebrafish hearts [15, 23, 25–28]. In this study, four CM populations were identified on the basis of the expression of *sorbs2b, ttn.2, ryr2b*, *myh7ba,* and *ldb3a* (**Fig. 1C, D** and **Supplementary Figure 1C, D**). To specifically study CMs in the regenerating heart, a second level clustering analysis was performed on the four designated CM clusters (clusters 0, 2, 6, and 7) (**Supplemental Figure 2A**). From this analysis, a total of eight distinct CM populations were identified (CM0-CM7) (**Fig. 1E-G**, **Supplementary Figure S2B-D, and Supplementary Table T2**). The analysis of genes enriched in the eight CM clusters revealed distinct biological functions based the distinct Gene Ontology (GO) terms for biological processes (**Fig. 1H, I**). A CM population involved in neonatal mouse CM cytoskeletal remodeling and injury response was classified and based on expression of *Ankrd1* and *Xirp2* [21]. In the adult zebrafish, CM7 is a population that shares similar gene expression profiles that include the *ankrd1a* and *xirp2a* genes (**Fig. 1F, G**). CM1 shares a high proportion of genes with CM7, but also includes transcripts involved in cell-cell contacts, sarcomere organization, and cardiac stress, highlighting that remodeling and dedifferentiation occur in the CM1 population (**Supplementary Table T2**). This highlights that remodeling and dedifferentiation are occurring in the CM1 population. Further evidence of CM dedifferentiation was noted by the expression of the *fos* and *jund* genes in the CM1 population (**Fig. 1H**, **Supplementary Figure S2C, D**). The GO biological processes associated with CM1 cluster included sarcomeres, myofibril assembly, and actomyosin organization, with the expression of *tcap, csrp3, and myoz2b* (**Supplementary Table T3**). CM0 exhibited the highest expression of genes involved in cardiac muscle contraction that includes *mylk4b, sorbs2b, nav2b,* and *chrm2A*. GO term designations confirmed that the biological processes associated with CM0 included genes involved in cardiac conduction and heart contraction (**Fig. 1H, I** and **Supplementary Table T3**). These attributes suggest that CM0 are spared CMs that function to maintain cardiac contraction and activity. Similarly, CM3 presented increased higher expression of cardiac contraction genes and represented another mature CM population (**Fig. 1H, I** and **Supplementary Table T3**). With respect to CMs in defined metabolic states, CM5 expresses genes involved in oxidative phosphorylation processes and likely represents homeostatic CMs (**Fig. 1H, I** and **Supplementary Table T3**). In contrast, CM2 expressed genes involved in insulin signaling and glucose metabolism (**Fig. 1F-I** and Supplementary Table T3). Such metabolic profiles in CMs have been described with glycolysis indicative of CMs linked to proliferation [15, 17]. Analysis of the CM4 gene expression profile revealed that cytoplasmic translation, and cellular proliferation are active processes in this population. The expression of the mitotic genes *ccnd2, skia, gli3* and *plk2b* supported the designation of CM4 as the proliferative population (**Fig. 1E-I**, and **Supplementary Table T3**). Finally, CM6 expressed genes associated with cell junction organization and heart development. CM6 likely represents a CM population that is undergoing remodeling (**Fig. 1H, I** and **Supplementary Table T3**).

**Figure 1.**
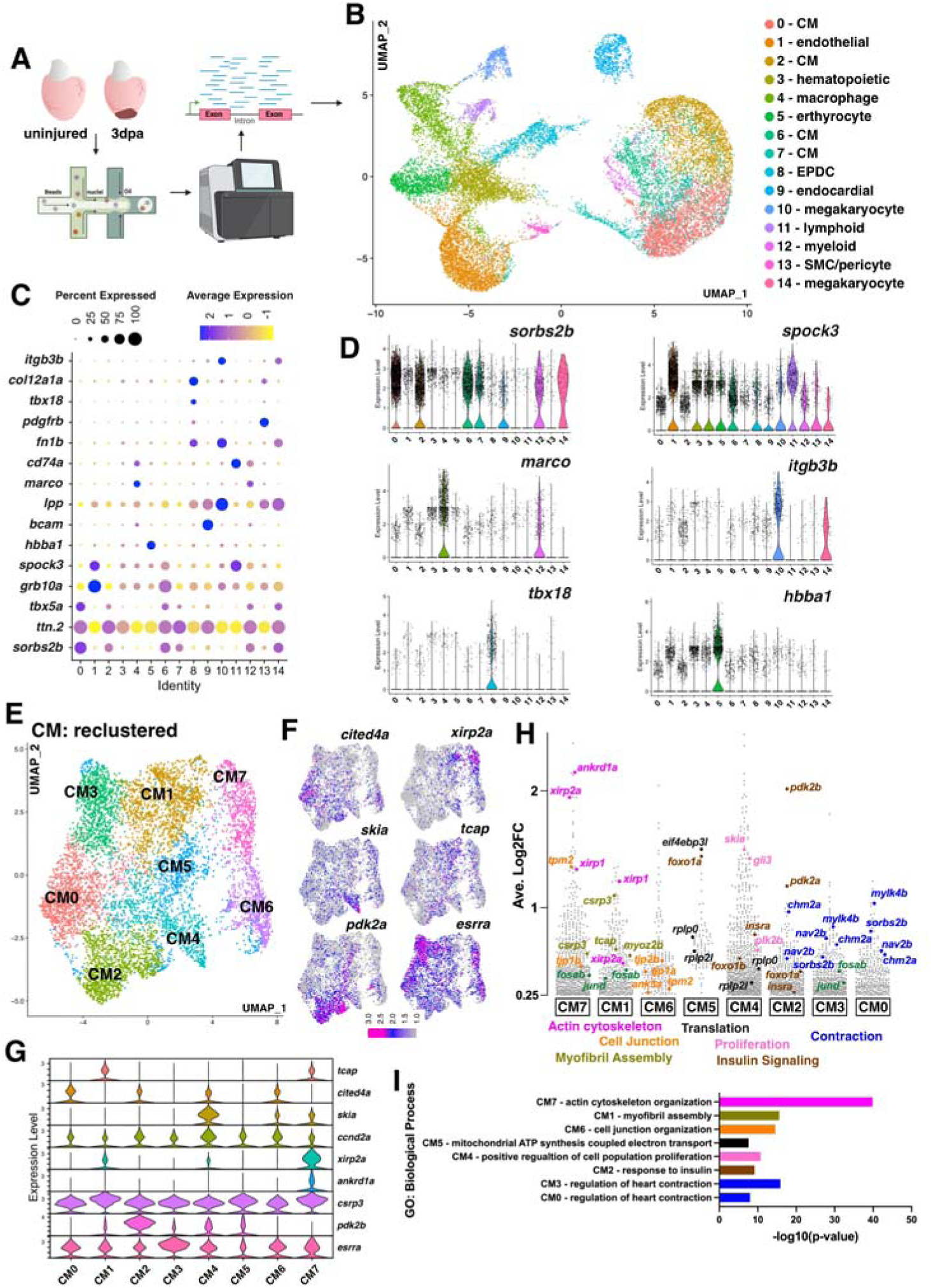
snRNA-seq of the regenerating adult zebrafish heart. **(A)** Overview of single nuclei RNA sequencing of zebrafish ventricles. **(B)** UMAP visualization of nuclei from snRNA-seq of uninjured and 3dpa hearts. Clusters are color coded with cell annotation. **(C)** Dot plot showing differentially expressed marker genes per cluster 0-14. **(D)** Violin plots showing expression levels of key cell type markers. **(E)** UMAP visualization of second level reclustered CM colored by cluster. **(F)** Feature plots visualizing distinct gene expression in CM clusters. **(G)** Violin plots showing expression levels of key markers across CM clusters. **(H)** Gene expression plot by average log2 fold change in each cluster. Each dot represents a single gene. Genes with functions related to actin cytoskeleton organization (fuchsia), cell junction rearrangement (orange), myofibril assembly (olive green), translation (black), proliferation (light pink), insulin signaling (brown), and contraction (blue). **(I)** Plot of top GO terms enriched for each cluster identified via ToppFUN bioinformatic tool.

We were intrigued by the CM populations defined by high expression of genes involved in CM maturation in the CM3 and CM0 populations. Specifically, *estrogen related receptor alpha* (*esrra*), *leucine rich repeat containing 10* (*lrrc10*), and *Cbp/p300 interacting transactivator 4a* (*cited4a*) are associated with CM maturation and cardiac remodeling (**Fig. 1F** and **Supplementary Table T3**) [29–33]. Lrrc10 was recently reported to be critical for CM maturation in regenerating zebrafish and mouse hearts [14, 34]. *ERRa* (*esrra* mouse ortholog) is a member of the steroid hormone nuclear receptor family and is reported to play a critical role in mouse CM maturation [29, 31]. To determine whether these *in silico* clusters represent distinct CM populations in the injured heart, RNA *in situ* hybridization was performed to determine the spatial expression of *cited4a, esrra* and *titin-cap* (*tcap)* at 3 dpa. *cited4a* was detected throughout the myocardium, but its expression was greater in CMs further from the border zone (**Fig. 2A- H**). The expression of *esrra* (CM3) was also greater in CMs distal to the plane of amputation (**Fig. 2A-D**). *tcap,* encoding a protein involved in sarcomere assembly, is a CM1-enriched gene that appears throughout the myocardium, but its expression is highest in the border zone (**Fig. 2G, H**). Fluorescence intensity measurements within multiple regions of the heart indicated that *cited4a* is positively correlated with *esrra* expression and negatively correlated with *tcap* (**Supplementary Figure S3A, B**). The proliferation genes, *prc1b*, *foxm1* and *g2e3* are specifically expressed in border zone CMs and have been used as markers for CM cell division [35]. From double *in situ* hybridization studies, *cited4a* wass only weakly expressed in proliferating *prc1b*, *foxm1* and *g2e3* positive CMs (**Supplemental Fig. S3C, D**). Moreover, the number of *cited4a+* CMs increased further distal from the border zone, in contrast to the presence of *prc1b* (**Supplemental Fig. S3E).** Taken together, the distinct CM populations identified from the snRNA-seq analysis are present *in vivo* and highlight CM heterogeneity in the regenerating zebrafish heart.

**Figure 2.**
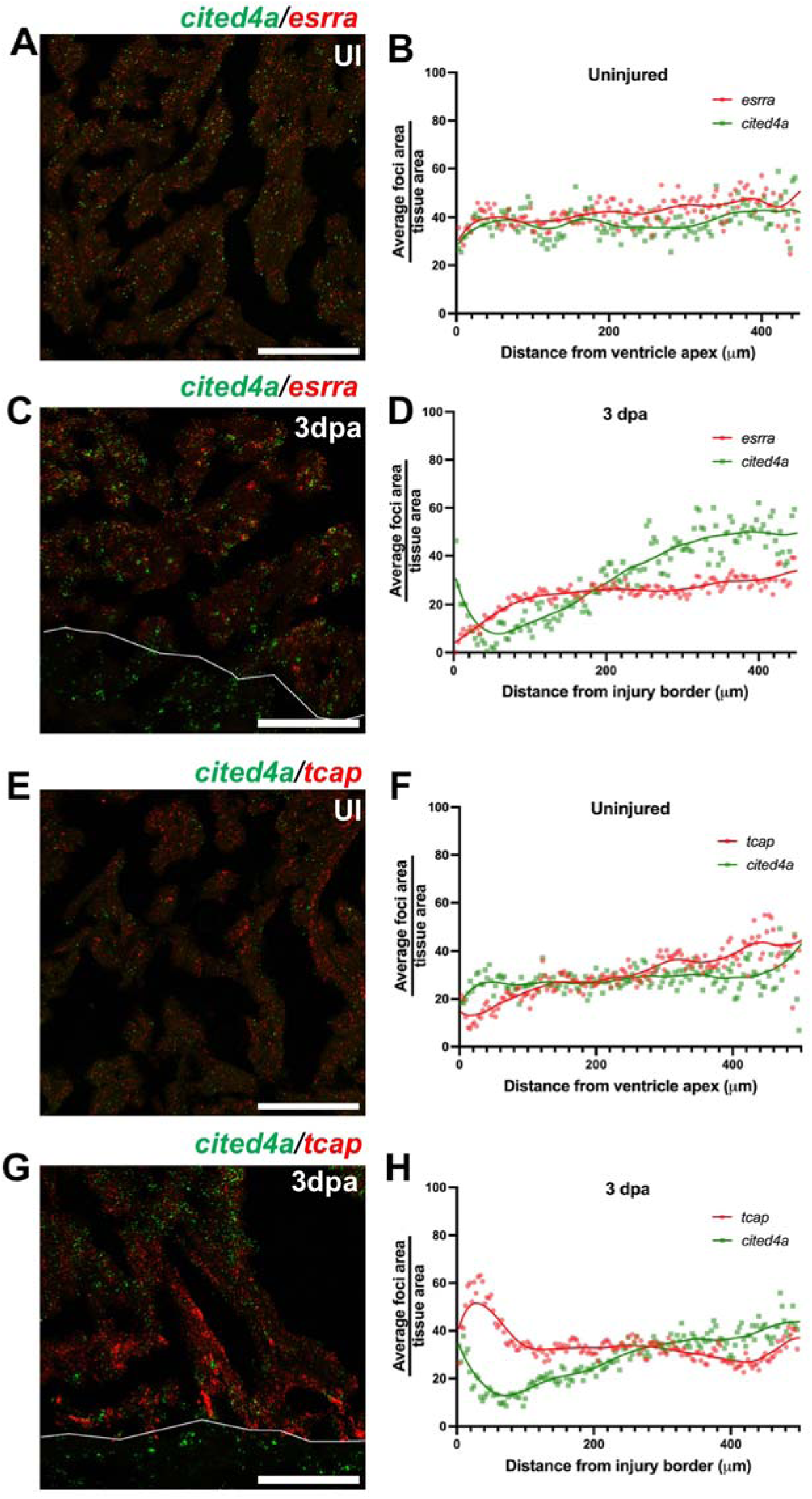
*cited4a, esrra* and *tcap* expression in CM populations in the heart. **(A-D)** Representative image of *cited4a* (green) and *esrra* (red) transcripts detected by RNAscope and quantification of signal foci/tissue area in relation to distance from the border zone in an uninjured (UI) ventricle (**A, B**) and injured (3dpa) (**C, D**). Representative image of *cited4a* (green) and *tcap* (red) transcripts detected by RNAscope and quantification of signal foci/tissue area in relation to distance from the border zone in an uninjured (**E, F**) and in 3 dpa ventricle (**G, H**). Scale bar: 100µm

The coexpression of transcriptional regulators *cited4a* and *esrra* in CM3 and CM0, suggests that *cited4a* may play an important role in CM maturation during regeneration. Interest in *Cited4* has been focused on its cardioprotective role against adverse remodeling after transverse aortic constriction in mice [33]. CITED4 induces healthy hypertrophy to protect the heart against detrimental remodeling in response to exercise and ischemic injury in mammals. *Cited4* is the least studied of the mammalian *Cited* genes and unlike *Cited1* and *Cited2*, *Cited4* is not required for embryonic development in mice [36–39]. In adult zebrafish heart, *cited4a* expression is induced after ventricular resection (**Supplementary Figure S4**).

### Cited4a disruption promotes heart regeneration

*cited4a* mutant zebrafish were generated via CRISPR/Cas9 to determine the role of this gene in heart regeneration. (**Figure 3A, B**). Homozygous mutants develop normally and survive to adulthood at the expected Mendelian ratio. The adult *cited4a^pt38a/pt38a^* hearts show increased volume, but no other defects were noted (**Supplemental Figure S5**). To determine how the loss of *cited4a* impacts heart regeneration, we quantified scar resolution and the expression of CM dedifferentiation and proliferation markers after injury (**Figure 3C**). Scar and fibrotic tissue in the amputated heart is typically resolved approximately 30 days after ventricular resection [8]. We noted that there was a decrease in fibrotic scar tissue at 20 dpa in the *cited4a* mutant (**Figure 3D-G**), suggesting that the resolution of the scar was accelerated in the mutant. Studies have shown that a unique myosin isoform and early embryonic sarcomere protein, embryonic cardiac myosin heavy chain (embCMHC, monoclonal antibody N2.261), is expressed in the larval heart and is a structural marker of embryonic undifferentiated CMs at 1 dpf [19]. embCMHC levels progressively decrease as the heart matures, but expression is reactivated after injury at the border zone, and is confined to within 100 μm of the injury at 7 dpa, where the majority of mitotic CMs are located [18, 19, 40]. Consistent with published data we observed embCMHC^+^ cells located within 100 μm of the injury border in WT hearts at 7 dpa and reach a maximum distance of 79 μm from the injury border (**Figure 3H-J**). In *cited4a* mutant ventricles, we observed embCMHC+ cells as far as 152 μm from the injury border. This trend was consistent in other *cited4a* allele, *pt38b* (**Supplemental Figure S6**). The expansion of embCMHC expression was accompanied by an increase in the number of proliferating CMs at 7 dpa, as quantified by CM expression of PCNA, *prc1b*, and *foxm1* in the *cited4a* mutant ventricles compared with that in the WT ventricles (**Figure 3K-S**). Together, these results indicate that *cited4a* is induced after injury to maintain a distinct CM population in a mature state, and in its absence, the pool of dedifferentiating and proliferating CMs increases, thereby accelerating the resolution of fibrotic scar tissue in the injured zebrafish heart. To identify pathways affected by *cited4a* loss in regenerating hearts, bulk RNA-seq of WT and *cited4a* mutant ventricles at 3 dpa was conducted (**Figure 4A and Supplementary Table T4**). Gene enrichment and functional annotation analyses of decreased transcripts revealed an overrepresentation of GO terms related to cardiac muscle contraction, fatty acid metabolic processes, and mitochondrial structure and function, suggesting decreased cardiac activity (**Figure 4B** and **Supplementary Table T4**). In addition, analysis of transcription factor perturbation followed by gene expression revealed significant overlap of genes whose expression decreased in the *cited4a* mutant with Esrra and Esrrg deficiency, and with Gata4 mouse knockout and gene ablation (**Figure 4B** and **Supplementary Table T4**).

**Figure 3.**
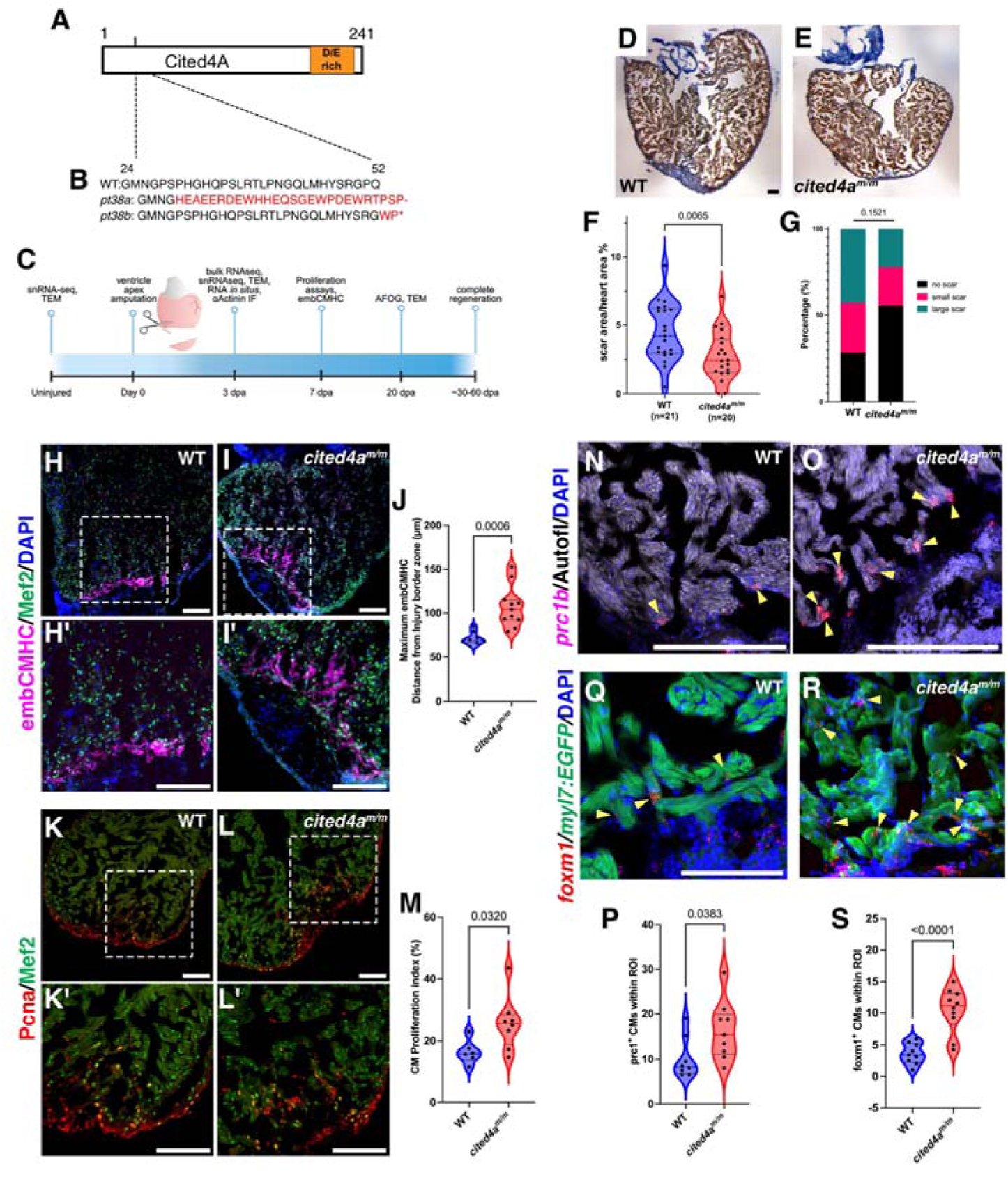
Cited4a limits CM dedifferentiation and proliferation in heart regeneration. **(A)** Schematic of Cited4a protein with glutamic and aspartic acid(D/E)-rich domain. (**B**) Predicted amino acid sequence of *cited4a* alleles. Altered sequence in red. **(C)** Timeline of injury and experiments. dpa, days post amputation. AFOG staining of heart sections at 20dpa to label fibrotic tissue (fibrin in red, collagen in blue) and muscle (orange-brown) in WT (**D**) and *cited4a^m/m^* (**E**). **(F)** Quantification of scar area as fibrotic area (μm^2^) divided by ventricle area (μm^2^). **(G)** Qualitative scoring of scar persistence at 20dpa: 0, no visible scar; 1, small amount of fibrosis; 2, medium to large amount of fibrosis analyzed by Fisher’s exact test. Immunostaining of embCMHC (magenta) to mark dedifferentiating CMs and Mef2a/c (green) to mark all CMs with DAPI (blue) in 3 dpa WT ventricles (**H, H’**) and *cited4a^m/m^* (**I, I’**). **(J)** Graph showing maximum distance of embCMHC staining from injury border zone in WT and *cited4a^m/m^*hearts. Immunostaining of Pcna (red) to mark proliferating CMs and Mef2 (green) to mark CMs in 3 dpa WT (**K, K’**) and *cited4a^m/m^* (**L, L’**) ventricles. **(M)** Quantification of CMs proliferation index (Pcna+;Mef2+/Total Mef2c+). *prc1b* (red) transcripts detected at the injury border zone by RNAscope with DAPI and autofluorescent myocardial tissue in 3dpa WT (**N**) and *cited4a^m/m^* (**O**) ventricles. **(P)** Quantification of proliferating *prc1b*+ nuclei in the myocardium. *foxm1* (red) transcripts detected at the injury border zone by RNAscope with *myl7:EGFP* to mark CMs and DAPI in 3dpa WT (**Q**) and *cited4a^m/m^* (**R**) ventricles. **(S)** Quantification of *foxm1+* CMs at the injury border zone. Scale bar: 100µm. For statistical analyses, Unpaired t-test were used unless indicated.

**Figure 4.**
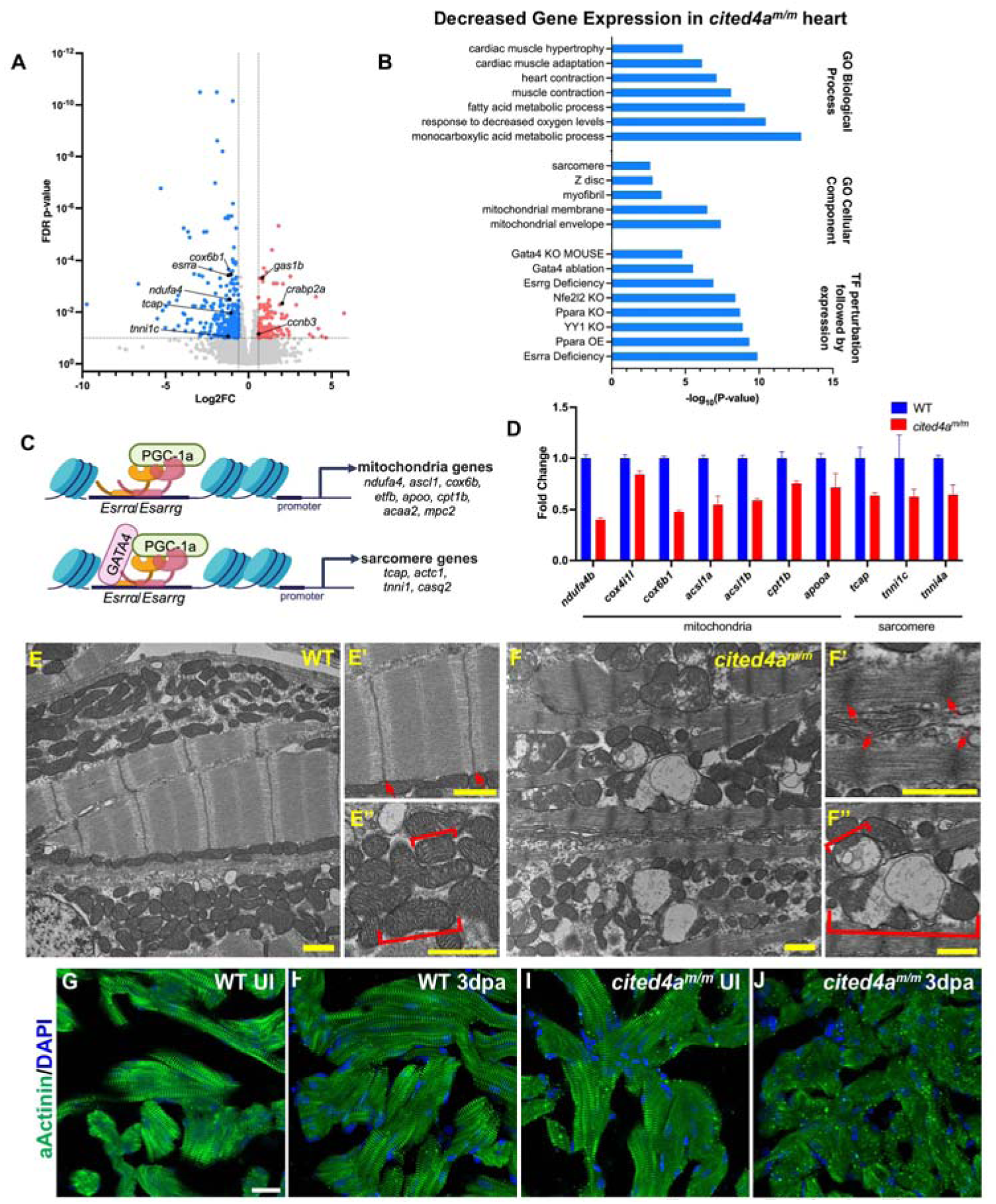
*cited4a* deficient regenerating hearts show altered mitochondrial and sarcomere structures. **(A)** Volcano plot of RNA-seq from *cited4a* mutant ventricles at 3dpa vs WT. Significantly decreased transcripts (blue), significantly increased transcripts (red), transcripts with no significant change (grey). **(B)** Biological process GO-terms, Cellular Component GO-terms and Transcription factor (TF) perturbation-associated gene expression signatures with significant overlap associated with significantly decreased genes in *cited4a^m/m^* mutant heart at 3dpa vs WT. **(C)** *Esrr*α*/Esrr*γ regulate CM maturation via induction of mitochondria and sarcomere gene expression. Designed with Biorender. **(D)** Representative qPCR validation of decreased mitochondria and sarcomere genes from bulk RNA-seq. Error bars represent standard deviation of technical replicates. Experiment was performed 3 times, showing consistent trends. **(E-E’’)** TEM of a WT 3dpa ventricle showing clear intact sarcomere structures (M-band, A-band, I-band, and z-disc) and mitochondria with densely packed cristae. (**F-F’’**) TEM image of a *cited4a^m/m^*mutant 3dpa heart showing a thick z-disk and dysmorphic mitochondria with fragmented cristae. Arrows indicate sarcomere z-disc. Bracket indicates a single mitochondrion. **(G-J)** Immunostaining of α-actinin to mark sarcomere z-disk and DAPI in uninjured (UI) WT (**G**), injured (3dpa) WT (**H**), UI *cited4a^m/m^* (**I**) and injured (3dpa) *cited4a^m/m^*(**J**) ventricles. Yellow scale bar:1µm. White scale bar: 20µm.

In the *cited4a* mutant heart, *esrra* expression was significantly decreased at 3 dpa (**Figure 4A**). ESRRα and ESRRγ, are the predominant ESRR isoforms in the mammalian heart and are highly expressed specifically in ventricular CMs [30] [31, 41]. ESRRα and ESRRγ regulate the expression of genes involved in metabolism and energy homeostasis via interaction with the transcriptional coactivator PGC-1α [42–45]. Moreover, it was recently shown that ESRRα/γ are necessary for the maturation of both mitochondria and sarcomere structure/function during postnatal cardiac development in mice [29, 31, 41]. ESRR⍰/γ form a complex with cardiogenic factor, GATA4, at CM-specific enhancer sites to activate the transcription of CM structural genes, whereas the regulation of metabolic gene occurs independently of GATA4 (**Figure 4C**) [29]. A significant number of genes whose expression decreased in the *cited4a* mutant were also downregulated in Esrrα [42] and Esrrγ [46] deficient mice (**Figure 4B**). Many of the genes whose expression were decreased are sarcomere- or mitochondria and are proposed targets of ESRRα/γ (**Supplemental Table T5**). Quantitative PCR (qPCR) was performed, and mitochondria and sarcomere genes identified from bulk RNA-seq confirmed decreased expression in the *cited4a* mutant hearts after ventricular resection (**Figure 4D**). To determine whether sarcomere structures and mitochondria are affected in CMs at 3 dpa as observed from the bulk RNA-seq and qPCR results, transmission electron microscopy (TEM) was performed. Like mature mammalian CMs, adult zebrafish CMs have ordered sarcomeres made up of actin and myosin filaments with well-defined z-lines and are surrounded by densely packed mitochondria (**Figure 4E** and **Supplementary Figure S7A**). Upon injury, sarcomeres near the border zone become disorganized and immature-like mitochondria can be observed (**Supplementary Figure S7C**) [13, 15, 16]. The mitochondrial structure in uninjured *cited4a* mutants was comparable to that in WT, but had a smaller area, perimeter, and more circular mitochondria (**Supplemental Figure S7D, I-K**). However, after injury at 3 dpa, we observed some striking mitochondrial defects in the *cited4a* mutant that were reminiscent of those in the hearts ESRRα/γ-deficient mice (**Figure 4F**, and **Supplemental Figure S7D, F**). At 3 dpa *cited4a* mutants CMs presented dysmorphic mitochondria with a rounded, swollen morphology and loosely packed or missing cristae were frequently observed (**Figure 4F** and **Supplemental Figure S7D, F**). In addition, blurring and thickening of the z-line in the *cited4a* mutant hearts after injury were also noted (**Figure 4F**, and **Supplementary Figure S7D, F**). Interestingly, CMs with irregular mitochondria were observed both at the border and in the remote zone in the mutant (**Supplemental Figure S7D, F)**. WT 3 dpa ventricles presented occasional and less severe mitochondrial morphology at the border zone, which is consistent with what has been described previously for border zone CMs and had a normal mitochondrial structure in the remote zone (**Supplementary Figure S7C, E**) [13, 15]. The *cited4a* mutant CM sarcomeres and mitochondria largely recovered by 20 dpa, whereas WT hearts still presented signs of injury (**Supplemental Figure S7G, H**).

### Cited4 is critical for CM maturation in hiPSC derived CMs

We next determined whether α-actinin localized in sarcomeres was altered in *cited4a* mutant hearts at 3 dpa. Uninjured WT and mutant ventricles displayed intact myofibrils/sarcomeres (**Figure 4G, I**), however at 3 dpa, *cited4a* mutants presented patches of disrupted α-actinin localization, indicating damaged myofibrils and disorganized sarcomeres (**Figure 4H, J**). Together, our bulk RNA-seq results combined with TEM analysis suggest a role for *cited4a* in maintaining normal CM mitochondria and sarcomere structures following injury.

We next investigated the role of human *CITED4* in CM maturation to determine whether it has a similar role in hiPSC-derived CMs. siRNA targeting *CITED4* or scrambled control siRNA were introduced into *ACTN2-eGFP* hiPSCs on differentiation day 13 or day 18 (**Figure 5A**). qPCR analysis at two days post-siRNA treatment revealed *CITED4* knockdown (**Figure 5B**). Sarcomere length via *ACTN2-eGFP* transgene expression was markedly shorter in CMs treated with *CITED4* siRNA than in those treated with scramble control siRNA (**Figure 5C, D, E)**. To gauge CM maturation, we measured the expression of two pairs of sarcomere genes that undergo fetal to adult isoform transitions: the troponin isoforms *TNNI3* and *TNNI1*, and the cardiac myosin heavy chain isoforms *MYH7* and *MYH6. TNNI3* and *MYH7* are the predominant isoforms in adult human CMs, whereas *TNNI1* and *MYH6* are the highest in the fetal heart [47, 48]. Following treatment with *CITED4* siRNA, the expression of the fetal isoforms was greater than that in the control CMs (**Figure 5G**), indicating that the CMs were less mature following *CITED4* knockdown. These results suggest that *CITED4* plays a pivotal role in promoting the maturation of hiPSC-derived CMs.

**Figure 5.**
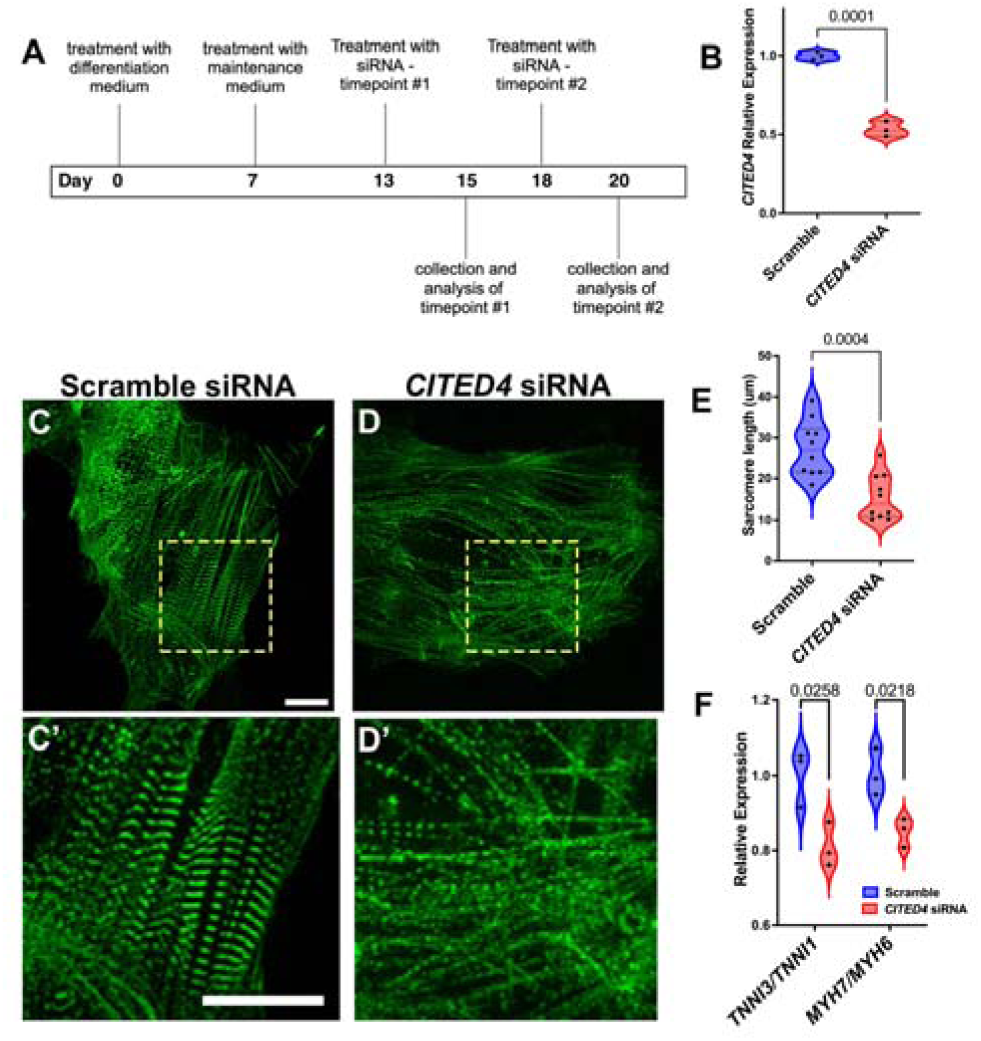
CITED4 is required for human iPSC derived CM maturation. **(A)** Experimental timeline. (**B**) qPCR analysis for *CITED4* expression in hiPSC-CMs following treatment with *CITED4* siRNA or scramble control siRNA ACTN2-eGFP hiPSC-CMs following treatment with scramble control (**C, C’**) of CITED4 (**D, D’**). Quantification of sarcomere length in hiPSC-CMs at Day 20 (**E**) following treatment with *CITED4* siRNA or scramble control siRNA. Gene expression ratios between *TNNI3/TNNI* and *MYH7/MYH6* measured by qPCR in hiPSC-CMs following treatment with *CITED4* siRNA or scramble control siRNA (**F**). Scale bar: 20 µm. Statistical analyses were by Unpaired t test.

### snRNA-seq of Cited4a mutant hearts reveals a distinct CM cellular state

We next performed snRNA-seq on *cited4a* mutant adult hearts before injury and at 3 dpa to determine the composition of CMs during regeneration in the absence of *cited4a*. In total we obtained 26,793 nuclei from uninjured and 3 dpa hearts, and 17 clusters were identified (**Figure 6A, B, Supplementary Figure S8,** and **Supplementary Table T6**). The contribution of cells from each of the conditions was uniform, suggesting a minimal batch effect (**Supplementary Figure S8**). The Harmony integration algorithm was employed for second-level clustering of the CMs, and to integrate the CMs between the various conditions. Harmony integration resulted in 14 distinct CM (CM0-13) clusters of which some CMs were common to WT and *cited4a* mutant hearts, but a large majority presented distinct states (**Figure 6C, D**, and **Supplementary Table T7**). The plots of the expression of genes that define injured (*xirp2a*), mature (*esrra* & *cited4a*), dedifferentiated (*tcap & jund*) and proliferating (*skia*) CMs revealed similar clusters that were previously identified via analysis of only WT CMs (**Figure 6E**, and **Supplementary Figure S9A-C**). The enriched genes in each CM cluster were used for gene expression profiling and GO term analysis, which resulted in distinct functional classes (**Figure 6F**, and **Supplementary Table T8**). Cardiac conduction and contraction genes were expressed in CM0, CM2 and CM8, indicating that these CMs are the most mature CMs (**Figure 6F, and Supplementary Figure S9D**). The injury response genes, *xirp2a,* and *xirp1* were expressed at the highest levels in CM4, CM5 and CM7. Notably, dedifferentiation genes, the AP-1 family of transcription factors, and *tcap* were enriched in CM8, CM5 and CM4. These three CM populations were predominately observed in the WT background and not in the *cited4a* mutant hearts (**Figure 6F**). According to the snRNA-seq data, *jund,* a member of the AP-1 family of transcription factors, is expressed at lower levels in *cited4a* mutant CMs. Recent work has revealed that AP-1 factors are activated in cardiac injury to induce the expression of the F-box protein involved in sarcomere protein degradation [18, 49]. We confirmed via *in situ* hybridization that *jund* expression is decreased in *cited4a* mutant hearts at 3dpa (**Figure 6G**). In addition, *esrra* was also reduced in the mutant hearts further supporting the lack of CM maturation in the *cited4a* mutant hearts (**Figure 6G**). A comparison of genes enriched in CM0 vs CM2, suggested that the expression of cardiac contraction and conduction genes was decreased in CM0 (**Supplementary Figure S9E**). This finding suggests that CM0, which is a population derived from *cited4a* mutant hearts is not as mature as the WT CM2. Thus, the CMs from *cited4a* mutant hearts appear to be immature when compared to WT hearts. Taken together, our analysis suggests that *cited4a* mutant CMs are immature when compared to CMs from adult WT and in a cellular state that can support survival and regeneration.

**Figure 6.**
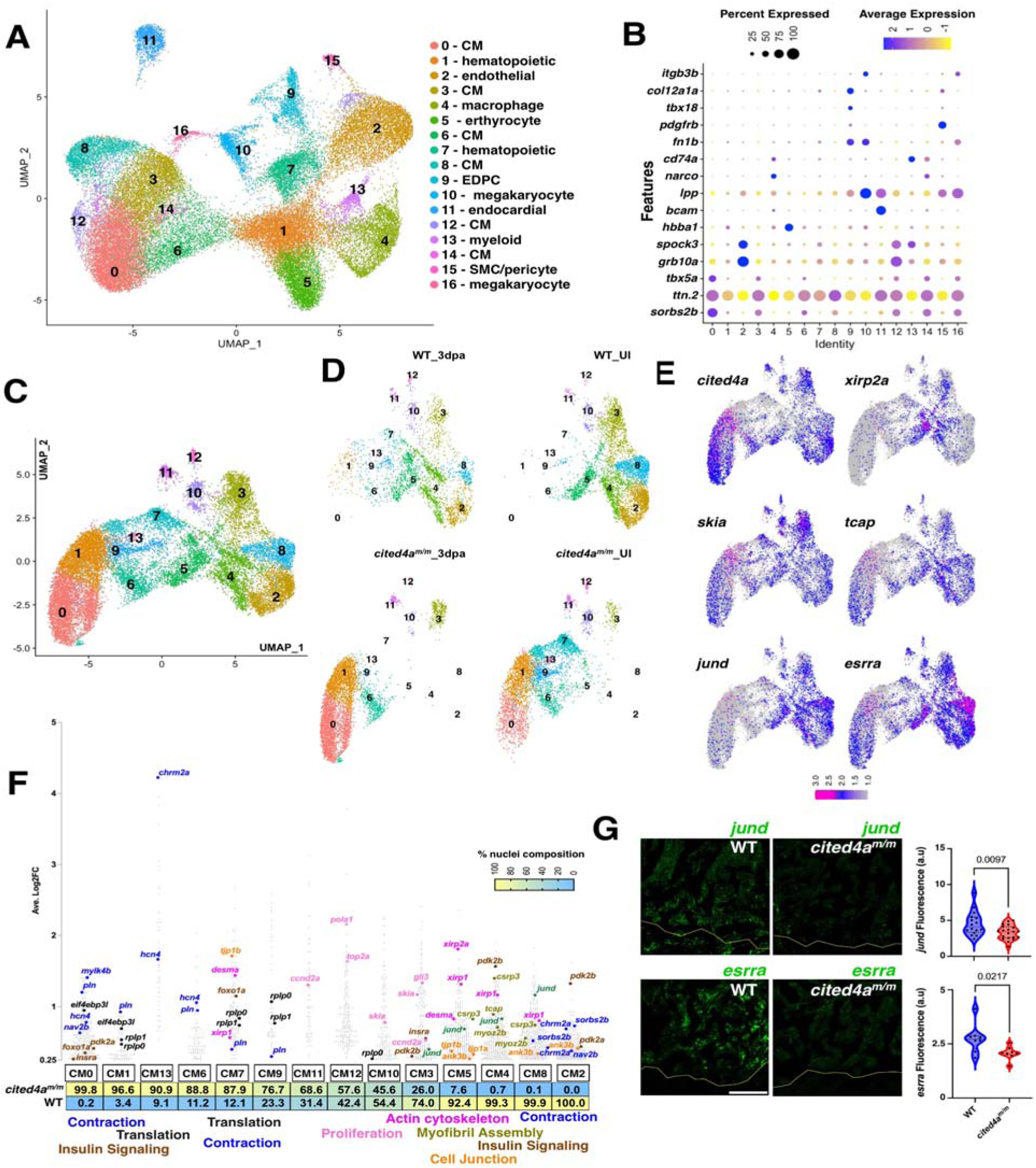
snRNA-seq of *cited4a* deficient hearts. **(A)** UMAP visualization of sequenced nuclei from all conditions (WT UI, WT 3 dpa, *cited4a^m/m^* UI, *cited4a^m/m^* 3 dpa) colored by cluster and labeled with cell type. **(B)** Dot plot showing differentially expressed marker genes per cluster. UMAP visualization of reclustered CM **(C)** and split into each condition and genotype **(D)**. **(E)** Feature plot of gene expression in CM clusters. **(F)** Gene expression plot by average log2 fold change in each CM cluster. Each dot represents a single gene, with percentage of nuclei composition for each genotype listed below. **(G)** Representative images of *jund* and *esrra* transcripts detected by RNAscope accompanied with graphs showing fluorescent intensity measurements in WT and *cited4a^m/m^* ventricles at 3 dpa. Unpaired t-test was used to determine significance with p-values indicated. Yellow line indicates site of injury. Fluorescent intensity of RNAscope in (a.u.=arbitary units). Scale bar: 100µm.

## Discussion

To investigate the transcriptional landscape of the regenerating zebrafish heart, we utilized snRNA-seq of uninjured and regenerating zebrafish ventricles. This snRNA-seq dataset provides a detailed, cell-type resolved picture of the zebrafish heart during homeostasis and regeneration and identifies new markers for distinct injury-responsive CM populations. Distinct CM populations that respond differently to injury and resemble those identified in regenerating neonatal mouse hearts have been identified. scRNA-seq of CMs from neonatal mouse hearts during regeneration identified five distinct CM populations, including one showing activation of an injury and stress response with cytoskeletal remodeling, similar to the CM7 and CM1 populations [21]. We were intrigued by maturation-associated genes, including *esrra*, expressed in CM0 and CM3. Both clusters show an overrepresentation of GO-terms associated with cardiac contraction. ESRRα and ESRRγ are critical for the maturation of CM mitochondria and for sarcomere structure and function during postnatal cardiac development in mice [31, 42–45]. The activation of this pathway in a subset of CMs during regeneration could serve as a cardioprotective mechanism. The heart must maintain function during regeneration, and dedifferentiating CMs have been shown to have decreased contraction and electrophysiological function. This is an underappreciated concept of the injury response where subpopulations are prevented from entering the cell cycle and instead increases CM maturation-associated processes, mitochondrial production and sarcomere integrity.

ESRRα and ESRRγ are activated by peroxisome proliferator-activated receptor γ coactivator-1α (PGC-1α) [42–45]. PGC-1α is regulated by p300, and PGC-1α can stimulate p300-dependent histone acetylation *in vitro* [50, 51]. *cited4a* could enable the *esrra* maturation pathway through its interaction with CBP/p300. Mammalian CITED4 is upregulated in response to cardiac stress such as endurance exercise or injury and can facilitate PGC-1α -stimulated histone acetylation by p300 when docked at ESRRα/γ nuclear receptors. Another model for the activation of the *esrra* maturation pathway via *cited4a* is inspired by a recent publication showing that the inhibition of p300 in cultured mouse myotubes led to an increased proportion of slow muscle fibers compared with fast muscle fibers [52]. Slow muscle fibers are rich in mitochondria, utilize oxidative metabolism, and are more resistant to fatigue. PGC-1α activates mitochondrial biogenesis and oxidative metabolism and can transform fast muscle fibers into slow muscle fibers [53], but this process can be inhibited by p300 [52]. In the heart, *cited4a* could sequester p300 from interacting with PGC-1α, allowing for activation of the *esrra* maturation pathway. It is also possible that *cited4a* activates the *esrra* maturation pathway through binding proteins other than CBP/p300. CITED1 and CITED2 have been shown to bind to other proteins such as SMAD3 [54], ISL1 [55], ERα [56],LHX2 [57], and PPARα [58], but it has yet to be demonstrated that CITED4 has other binding partners *in vivo*. Together, these data suggest that *cited4a* acts upstream of *esrra* in CMs during the early stages of zebrafish heart regeneration to maintain CM maturity. These findings suggest that the modulation of endogenous maturation programs could increase the regenerative potential of the heart and could be used as a therapeutic approach.

This study highlights a role for *cited4a* in the activation of CM maturation pathways as an injury response. We have shown that in the absence of *cited4a*, increased CM dedifferentiation and proliferation are observed. In addition, loss of *cited4a* leads to significant transcriptional changes in CMs during homeostatic conditions and after injury. Gene ontology analysis of CM clusters containing exclusively mutant nuclei revealed an enrichment of genes related to hypertrophic cardiomyopathy. In mice, *Cited4* is proposed to act as part of the physiological hypertrophy response program and induce remodeling to protect the heart against pathological hypertrophy, which involves ventricular dilation, reduced systolic function, and increased fibrosis. One characteristic feature of pathological cardiac hypertrophy is the re-expression of fetal genes [59]. The *cited4a* mutants presented increased expression of embCMHC and a transcriptional profile that suggested an irregular hypertrophic response. These observations suggest that *Cited4* function is conserved between mice and zebrafish during cardiac stress response. Furthermore, knockdown of *CITED4* in hiPSC-CMs led to increased expression of embryonic isoforms and shorter sarcomere length, suggesting a role in CM maturation. Together, these data suggest that *cited4a* acts upstream of *esrra* in CMs during the early stages of zebrafish heart regeneration to maintain CM maturation. These findings suggests that the modulation of endogenous maturation programs could increase the regenerative potential of the heart and could be used as a therapeutic approach.

## Materials and Methods

### Zebrafish lines

*cited4a* mutant alleles were created using CRISPR/Cas9 according to established protocols with sgRNA listed in **Supplementary Table T9** [60]. Genotyping was performed by Restriction Fragment Length Polymorphism assay [61], and the PCR product was digested using restriction enzyme DdeI (NEB, R0175S). Two independent alleles were established: *pt38a* (95bp deletion) and *pt38b* (19bp deletion). Both alleles contained deletions resulting in a frameshift and early stop codon and the *pt38a* allele was used for all experiments unless otherwise noted. The PCR primers used are listed in **Supplementary Table T9**.

### Zebrafish ventricle resection and tissue processing

Ventricle resection in adult zebrafish (6-12 months) was performed following established protocols [8]. Extracted hearts were washed in cold PBS. Unless otherwise stated, the hearts were incubated in 4% PFA overnight at 4°C. The hearts were then washed in PBS and transferred into a sucrose gradient (10%, 20%, and 30% sucrose in PBS) and stored in 30% sucrose at 4°C overnight. Hearts were mounted in Surgipath Cryo-Gel (Leica Biosystems #39475237), frozen at −80°C, and sectioned on a Leica CM1850 cryostat at 14µm. For IF staining unfixed hearts were mounted in OCT media (Scigen #4586), flash frozen on dry ice in an ethanol bath, and sectioned at 10 µm. The tissue was then fixed on the slide with 4% PFA.

### Zebrafish Immunofluorescence and RNAscope

For Mef2 and PCNA staining, slides with heart sections were rehydrated PBS-Tween (0.1% Tween in PBS) and permeabilized with 4% hydrogen peroxide in methanol for 1 hour. The slides were then incubated in IF blocking buffer (2% sheep serum, 1% DMSO, 0.2% TritonX- 100 in PBS) for 1 hour. Primary antibodies were diluted in blocking buffer, added to slides and incubated overnight at 4°C. Secondary antibodies diluted in blocking buffer were added and incubated for 2 hours at room temperature or overnight at 4°C. The slides were then incubated with DAPI for 5 minutes and sealed with Aqua Poly/Mount and a coverslip. For MEF2A+MEF2C, α-Actinin, and embCMHC, slides were rehydrated TBS-Triton (0.2% Triton X-100 in TBS) and permeabilized in 4% Triton X-100 in TBS for 1 hour. Slides were then incubated in IF blocking buffer (1% BSA, 5% sheep serum, 0.2% TritonX-100 in TBS) for 1 hour. Primary antibodies were diluted in blocking buffer, added to slides and incubated overnight at 4°C. Secondary antibodies diluted in blocking buffer were added and incubated for 2 hours at room temperature or overnight at 4°C. Slides were then incubated with DAPI for 5 minutes. See **Supplementary Table T9** for antibodies and vendor information.

RNAscope in situ hybridization was performed as previously described [62] on 14 µm tissue sections using the RNAscope Multiplex Fluorescent Detection Kit (ACD # PN323110) and following the protocol for RNAscope Multiplex Fluorescent Reagent Kit v2 (ACD). TSA Cy3, Cy5, and Fluorescein dyes (PerkinElmer # NEL760001KT) were used at a ratio of 1:750. RNAscope probes are listed in **Supplementary Table T9**. All IF and RNAscope images were acquired on a Zeiss 700 confocal microscope.

### Zebrafish Immunofluorescence and RNAscope quantification

The average and maximum distances of embCMHC-expressing cardiomyocytes was quantified at using ImageJ (FIJI). Ten regularly spaced lines were drawn perpendicular to the injury border zone, reaching the edge of the embCMHC expression. Four sections were quantified per individual heart and the average and maximum distances from the injury border zone were subsequently calculated for each individual section. Also, the average of average and maximum distances were measured and calculated for each heart, and plotted. A minimum of three biological replicates was performed for each experiment and experiments were repeated three times. Cardiomyocyte proliferation index (%) was calculated from the number of Mef2a/c+;Pcna+ double-positive cells divided by the total number of Mef2a/c+ cells. Four sections were quantified per individual heart and a minimum of three biological replicates used in each experiment and repeated three times. Protein regulator of cytokinesis 1b (*prc1b*)- expressing cardiomyocytes were quantified from confocal images of the ventricle BZ taken with a 40X water objective. Quantification was performed by counting the number of nuclei (DAPI) within autofluorescent myocardial tissue containing at least 3 RNAscope probe puncta. *prc1b* scoring was performed by an experimenter blinded to the genotype. Two sections were quantified per individual heart, and a minimum of four biological replicates used in each experiment and repeated two times.

RNAscope signals were quantified via multiple methods. The distance and number of *cited4a^+^* and *prc1b^+^* cells from the BZ were quantified according to previously established methods [35]. Transcript levels of *cited4a*, *tcap*, and *esrra* were collected as the mean intergraded density (signal normalized by the area) from ten ROIs within the tissue area on individual slices from four ventricles. Specifically, ten 50X50 pixel ROIs were randomly placed within the tissue area, and the mean intergraded density was measured for each channel using FIJI. The results were plotted as x-y coordinates to show expression correlation. To measure transcript levels (foci area) in relation to the distance from the ventricle apex or injury border zone, images were analyzed using Python (v. 3.9) with image analysis library Scikit-Image (v. 0.19.3) [63]. The line defining the edge of the tissue in healthy hearts, or the border of the injury was defined manually. The background in the images was defined using a median filter (radius 15 px) and subtracted from the images before the analysis. Foci were defined by setting a consistent signal threshold. As targeted RNAs are often detected in dense clusters, we present the results as the ratio between the area covered with foci and the total area of the tissue. For presentation, the data were normalized to values ranging from 0 to 100. Sarcomere length was measured in zebrafish via αActinin immunostaining. A line was drawn across a CM, perpendicular to the z-lines using ImageJ to measure the distance between the z-lines. Seven to ten individual sarcomere lengths were collected per cell.

### Zebrafish Histology

Acid fuchsin orange G (AFOG) staining was performed as previously described [8]. Images were taken with a Leica MZ16 microscope and a Q Imaging Retiga 1300 camera. The scar area was calculated as the fibrotic area divided by ventricle area. The area was measured via ImageJ, with 4 sections per individual heart. Semi-qualitative scoring of scar persistence at 20 dpa was performed following a modified version of published methods [35]: 0, no scar; 1, small amount of scar; 2, medium to large amount of scar. Scoring was performed by an experimenter blinded to the genotype.

### Zebrafish ventricle bulk qPCR and RNA-seq

Zebrafish ventricle bulk RNA-seq Hearts were extracted from WT and *pt38a* zebrafish at 3dpa and the atrium and bulbus arteriosus were manually removed. Total RNA was isolated using Trizol (ThermoFisher #15596018) and RNeasy micro kit (Qiagen #74004). For qPCR, eight ventricles were pooled per condition. Between 0.5-1μg of total RNA was used to generate cDNA via the SuperScript First-Strand Synthesis System (ThermoFisher # 11904-018). The mRNA expression levels were determined by quantitative PCR using a Power SYBR™ Green PCR Master Mix Kit (Applied Biosystems, 4368706) as previously described [64]. The expression values were normalized to those of *actb2* or *polr2d* mRNA and were represented as fold change. The qPCR primers used are listed in **Supplementary Table T9**. A representative example is shown with the mean and standard deviation of three technical replicates from one biological experiment. Three independent biological replicates were performed. For RNA-seq, 5-8 ventricles were pooled per condition and four biological replicates were used for bioinformatic analysis. mRNA libraries were constructed for paired-end sequencing at a depth of 40 million reads per sample. Library preparation and sequencing were performed at the Genomics Research Core at the University of Pittsburgh. The RN-seq reads were adaptor trimmed, mapped to the zebrafish genome and the reads were counted using standard procedures of FastQC, HiSat2 and featureCount packages. The R package DEseq2 was used to analyze differential gene expression (DGE) between *cited4a* mutant and WT hearts at 3 dpa. Enrichr was used to find overlapping genes from our DEGs and previously annotated gene sets from transcription factor perturbation experiments [65]. Bulk RNA-seq sequence files and featureCount expression counts are available at GEO under the accession number: GSE279843

### Zebrafish ventricle single nuclei RNA sequencing (snRNA-seq)

Hearts were extracted from uninjured and 3dpa from WT or *cited4a^pt38a^*zebrafish for nuclear isolation using the Chromium Nuclei Isolation Kit (10x Genomics #1000494). The atrium and bulbus arteriosus were manually removed. Four ventricles were pooled per condition. Library preparation was performed at Single Cell Core at the University of Pittsburgh (singlecell.pitt.edu). Sequencing was performed at the UPMC Genomics Center. A total of 10,000 nuclei were sequenced at a depth of 20,000 reads per cell. The raw sequence files are available at GEO under the accession number: GSE279845.

### Mapping and Bioinformatics analysis

Raw FASTQ files were processed and aligned via the Cell Ranger (v. 7.0.0) “count” utility with automatic chemistry detection, aligning to the *Danio reiro* genome (GRCz11.108) [66]. Processed counts were read into R (v4.2.0) using the Seurat package (v4.3.0). Nuclei were filtered for a minimum of 3 cells and 200 RNA features [67, 68]. The samples were filtered to exclude nuclei with abnormally high mitochondrial gene expression and RNA features to remove potential contamination and/or doublets. The samples were then integrated following the standard Seurat integration pipeline, as follows: First, the data were scaled and normalized separately for each sample, and the top variable features were calculated. Then, Principal Components Analysis (PCA) was subsequently performed, and the integration anchors between sample datasets were calculated via the top 2000 variable features and top 20 principal components identified using comprehensive Jackstraw and Elbow heuristics. The samples and calculated anchors were passed to the “IntegrateData” function, resulting in an integrated Seurat object. This integration procedure was performed three separate times: 1) Integration of the Wild-Type Uninjured (WT-UI) and Wild-Type 3dpa (WT-3dpa) samples. 2) Integration of the *cited4a* Mutant Uninjured (Mutant-UI) and *cited4a* Mutant 3dpa (Mutant-3dpa) samples. 3) Integration of the integrated Wild Type (WT-UI + WT-3dpa) and integrated Mutant (Mutant-UI + Mutant-3dpa) datasets. The integrated datasets were scaled and normalized, and the top Principal Components were calculated using the procedure described above. Clusters were generated using Seurat’s “FindNeighbors” and “FindClusters” functions using the top 15 identified principal components and a clustering resolution of 0.5. Uniform Manifold Approximation and Projection (UMAP) dimensionality reduction was performed utilizing the top 15 principal components. Cellranger files are available at GEO under the accession number: GSE279845

### Cell Type Annotation

The top positive marker genes for each cluster in the integrated datasets, were calculated using the FindAllMarkers function from Seurat. Genes were selected on the basis of a minimum expression percentage of 0.25 and a logFold-Change expression threshold of 0.25. The top 50 cluster-specific markers were utilized for manual cluster annotation.

### Differential Expression

Differentially expressed genes between conditions for each cluster were calculated using the FindMarkers function from Seurat. Genes were determined to be differentially expressed with a Bonferroni-adjusted p-value < 0.05 and | logFold-Change | > 0.25. The marker genes form each cluster was converted to human gene orthologs via gProfiler [69]. GO term analysis was performed on each cluster with the human gene orthologs via ToppFUN from the ToppGene Suite [70].

### Second-level Reclustering of Wild type and *cited4a* Mutant Cardiomyocytes

Annotated CMs from the integrated Wild type dataset were subset and reclustered following the procedure described above to achieve higher resolution clusters of CMs and investigate distinct levels of maturation. We then utilized Harmony integration to compare the expression levels of maturation markers between the integrated WT and *cited4a* Mutant CMs [71]. Having annotated the integrated WT+Mutant clusters as described above, we subset the annotated cardiomyocyte clusters and used Harmony to reintegrate the subset nuclei and generate a harmonized dimensionality reduction, to eliminate potential batch effects between experimental groups that were observed when the standard Seurat clustering procedure was used. Harmonized cell embeddings were used in lieu of PCA embeddings for the FindNeighbors, FindClusters and the RunUMAP clustering procedure described above. This resulted in a refined set of overlapping and distinct clusters between the Wild type and mutant nuclei suitable for further analysis and developmental inference.

### Zebrafish TEM

Ventricles were fixed in cold 2.5% glutaraldehyde in 0.01 M PBS. The samples were rinsed in PBS, post-fixed in 1% osmium tetroxide with 1% potassium ferricyanide for 1 hour, rinsed in PBS, dehydrated through a graded series of ethanol and propylene oxide and embedded in Epon resin (Poly/Bed® 812). Semi-thin (300 nm) sections were cut on a Leica Reichart Ultracut, stained with 0.5% Toluidine Blue in 1% sodium borate and examined under the light microscope. Ultra thin sections (65 nm) were stained with uranyl and one section was randomly selected for imaging on JEOL 1400 Plus transmission electron microscope (grant # 1S10RR016236-01 NIH for Simon Watkins) with a side mount AMT 2k digital camera (Advanced Microscopy Techniques, Danvers, MA). For the 3 dpa and 20 dpa samples, ventricles were bisected from the remote zone (RZ) and border zone (BZ), processed, and imaged separately. The BZ sample contained the injury site and adjacent tissue. Uninjured ventricles were bisected and only the sample containing the ventricle apex was processed and imaged. For uninjured and 20 dpa n=1; for 3 dpa, n=2.

Sarcomere and mitochondrial ultrastructures were evaluated via a modified semiquantitative scoring system utilized previously [72, 73]. For each sample, 10 nonadjacent regions at a primary magnification of 10000x-15000x were randomly selected for scoring according to a four-point scale describing the degree of ultrastructural damage. Scoring was performed by an experimenter blinded to the genotype. Mitochondria shape (area, perimeter, circularity) measurements were performed by tracing individual mitochondria in ImageJ (FIJI) and the measure function. For each sample, five non-adjacent regions at a primary magnification of 10000x-15000x were randomly selected for scoring.

### Monolayer cardiac differentiation from hiPSCs and maintenance

Monolayer cardiac differentiation was carried out following a published protocol (PMID: 26125590). Cardiac cells from monolayer differentiation cultures at day 15 and day 20 were washed twice with PBS and incubated with TrypLE Express (Life Technologies #A1217702) for 5 min at 37 °C. The cells were collected by centrifuge at 300 x g for 5 min, resuspended in 1 ml of B27 and filtered through a 40 μm filter (Corning #431750).

The WTC line with the ACTN2-eGFP transgene (Coriell # AICS-0075-085) was maintained in completely defined albumin-free E8 medium (DMEM/F12 with L-glutamine and HEPES, 64 μg/ml L-Ascorbic Acid-2-phosphate, 20 μg/ml insulin, 5 μg/ml transferrin, 14 ng/ml sodium selenite, 100 ng/ml FGF2, 2 ng/ml TGFb1) on Matrigel (Corning #CB40230A) coated tissue culture plates. The medium was changed daily and routinely passaged every three to four days using 0.5mM EDTA solution (Invitrogen #15575020).

### CITED4 RNAi in hiPSCs

The CITED4 gene was knocked down by transfection with small interfering RNA (siRNA) targeting CITED4 (Thermo Fisher #s225786) via Lipofectamine™ RNAiMAX (Invitrogen #13778). Cells were seeded in 2.5 ml of RPMI-B27 per well in 6-well plate with 50% confluence for transfection. On the day of transfection, 5 μl lipofectamine and 10 nM RNAi duplex were diluted in 250 μl of Opti-MEM® I Reduced Serum Medium (Gibco #31985-062) separately, before being mixed and incubated for 20 min. The mixture was then added to the culture medium and cells were incubated at 37 in a CO_2_ incubator for 6 hours. The medium was changed back to RPMI-B27 after 6 hours and gene knockdown was confirmed by real-time qPCR after 48 hours.

### Sarcomere Measurements in hiPSCs

For cell imaging, the resuspended cells were seeded onto coverslips and imaged under Leica TCS-SP8 confocal microscope. Sarcomere length was measured using via α-actin eGFP from 5 individual images in control and CITED4 knockdown cells separately.

### RNA extraction and qPCR in hiPSCs

RNA was extracted, and purified, and mRNA expression levels were determined via the same method used for zebrafish as described above except cDNA was synthesized using an iScript™ cDNA Synthesis Kit (BIO-RAD, 1708891). The primers used in this study are listed in **Supplementary Table T9**.

## Acknowledgements

Support for the research are from grants to M.T. (NIH 1R01HL156398), and D.Z. (F31 HL149148).

This research was supported in part by the University of Pittsburgh Center for Research Computing through the resources provided. Specifically, this work used the HTC cluster, which is supported by NIH award number S10OD028483. Additionally a grant from the Chan Zuckerberg Initiative DAF (number 2023-329680), an advised fund of Silicon Valley Community Foundation supported part of this study.

## Author Contributions

R.F.R., and M.T. initiated the project concept. R.F.R performed the snRNAseq and bulk RNA experiment. B.S., D.R. and M.T. analyzed the snRNAseq and bulk RNAseq data. R.F.R. generated zebrafish *cited4a* mutants. R.F.R., A.P., D.Z., performed RNA in situ and with K.K. analyzed the data. R.F.R, D.Z., A.P., performed heart regeneration studies and analyzed the data. W.F., and G.L., performed hiPS studies. R.F.R, D.S., and S.W., performed TEM imaging and analysis. R.F.R and M.T., analyzed all the data, prepared figures, and drafted the manuscript. R.F.R., B.S., W.F., G.L., D.R., A.P., D.Z., and M.T., edited the manuscript. All the authors have read the manuscript.

## Supplementary Figures

**Supplementary Figure S1.**
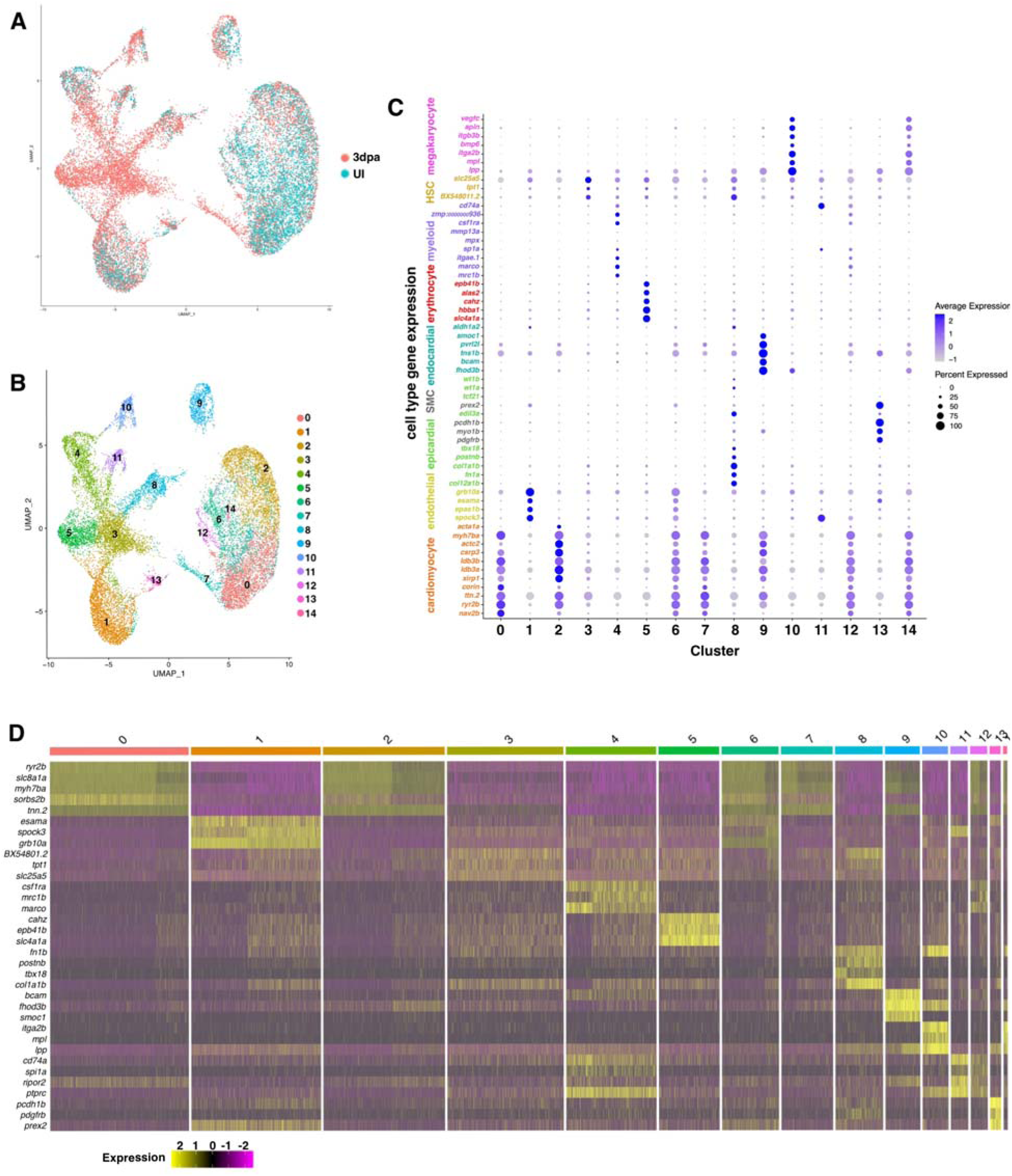
snRNA-seq of the regenerating adult zebrafish heart. UMAP visualization of nuclei from snRNA-seq of uninjured and 3dpa hearts split by stage (**A**) and showing the 15 clusters (**B**). Dot plot (**C**) and heatmap (**D**) gene expression profile of the 15 Clusters from the integrated uninjured and 3dpa heart.

**Supplementary Figure S2.**
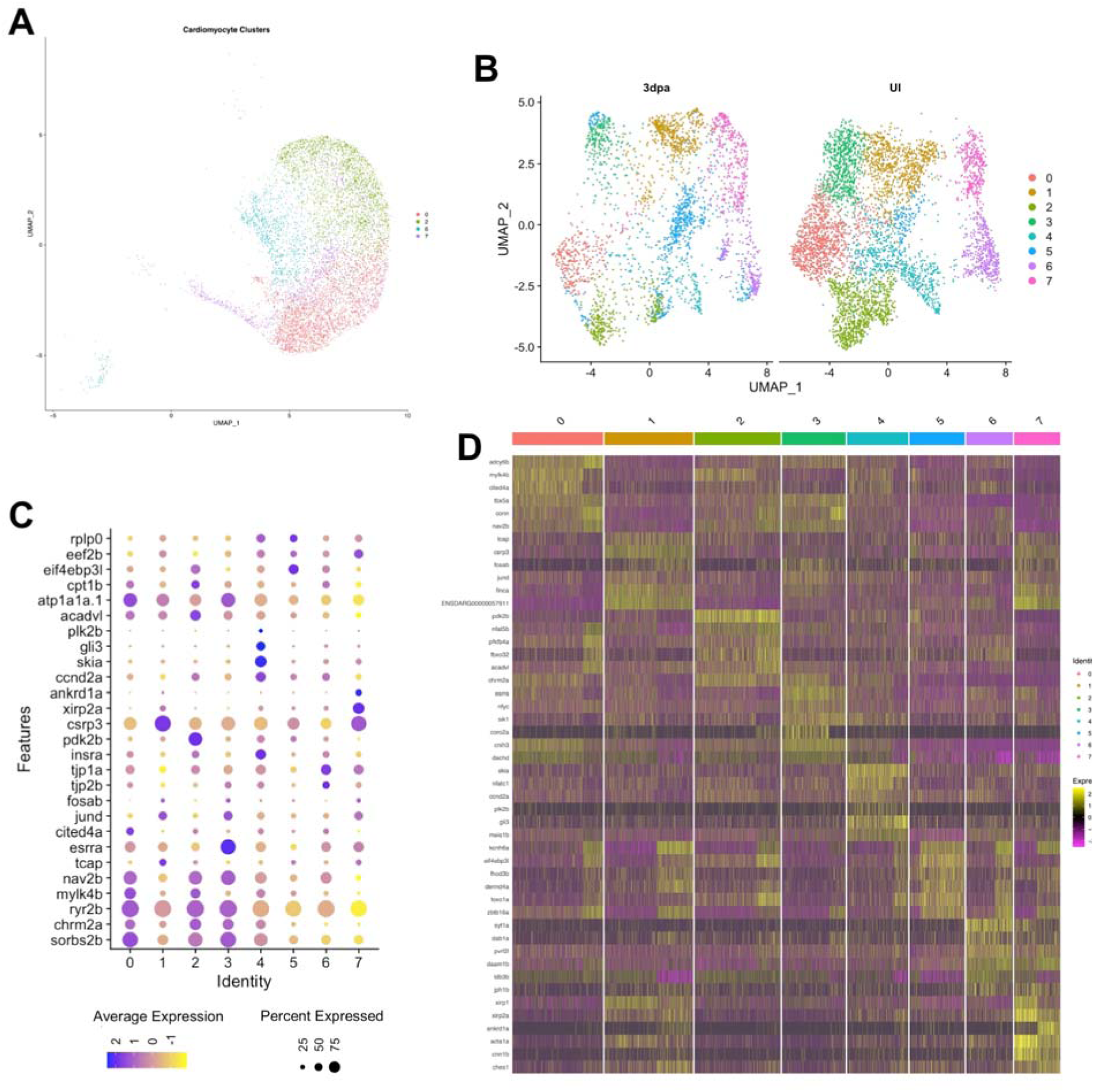
Second level analysis of CM from Uninjured and 3dpa hearts. **(A)** UMAP visualization of CM clusters. (**B**) UMAP visualization of second level reclustered CMs colored by cluster. Dot plot (**C**) and heatmap (**D**)showing differentially expressed marker genes from the reclustered CMs.

**Supplementary Figure S3.**
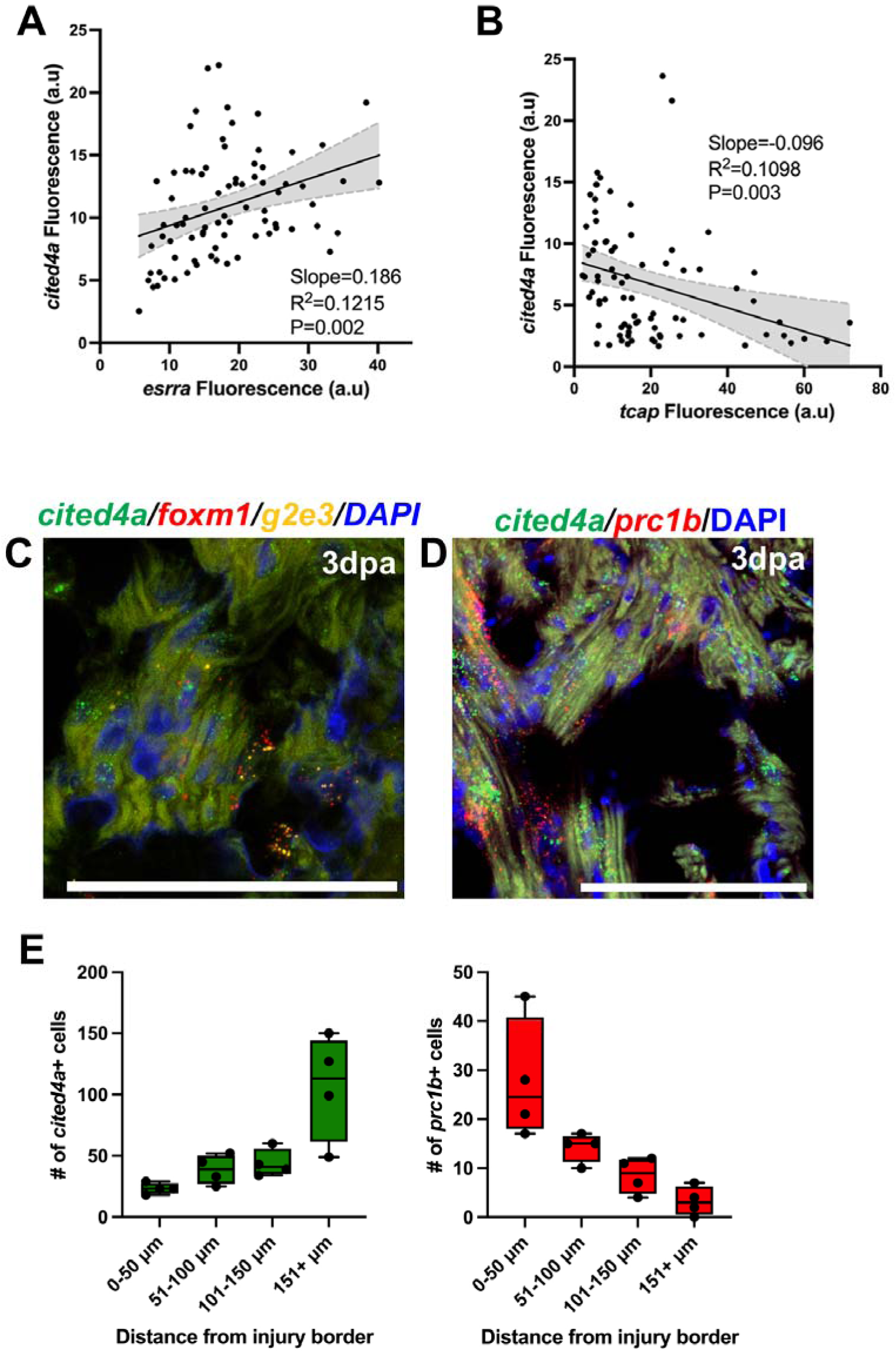
*cited4a* expression in injured heart. **(A, B)** *cited4a, esrra* and *tcap* RNAscope fluorescence correlation quantification from 3dpa hearts. *cited4a* CM expression is co-expressed with *esrra,* but not with *tcap*. (**C, D**) Double and triple RNAscope staining showing *cited4a* with proliferation marker genes, *foxm1*, *g2e3* and *prc1b*. (**E**) Quantification of *cited4a* and *prc1b* positive CMs in relation to distance from the injury border. Scale bar = 100µm.

**Supplementary Figure S4.**
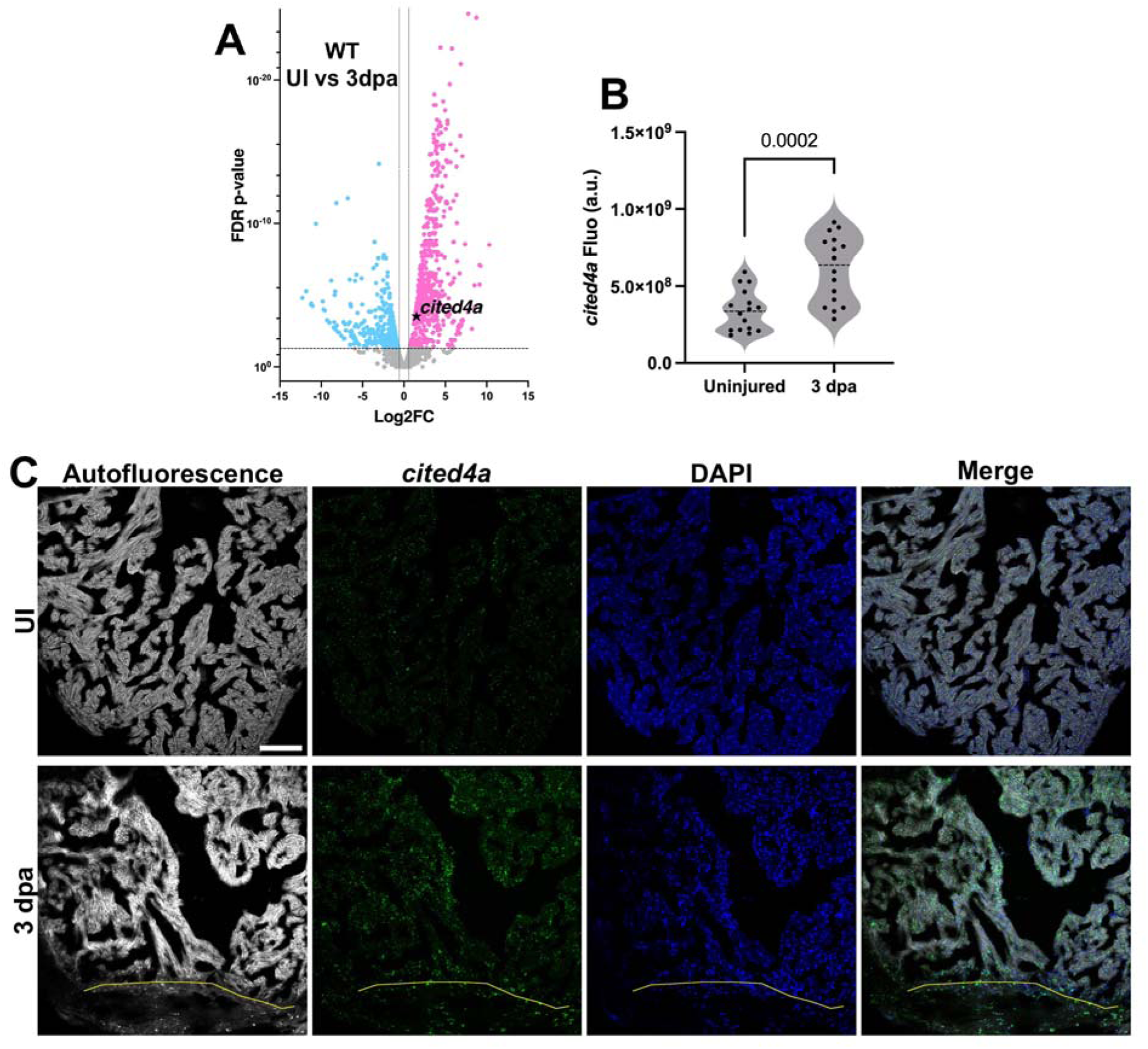
*cited4a* is induced in the heart after ventricular amputation. (**A**) Volcano plot of RNA-seq data showing *cited4a* expression is induced in the injured heart from *Zuppo et al*. 2023. (**B**) Quantification of RNAscope *cited4a* staining in the heart before and after injury. (**C**) Representative images of *cited4a* RNAscope in situ hybridization. Scale bar = 100µm.

**Supplementary Figure S5.**
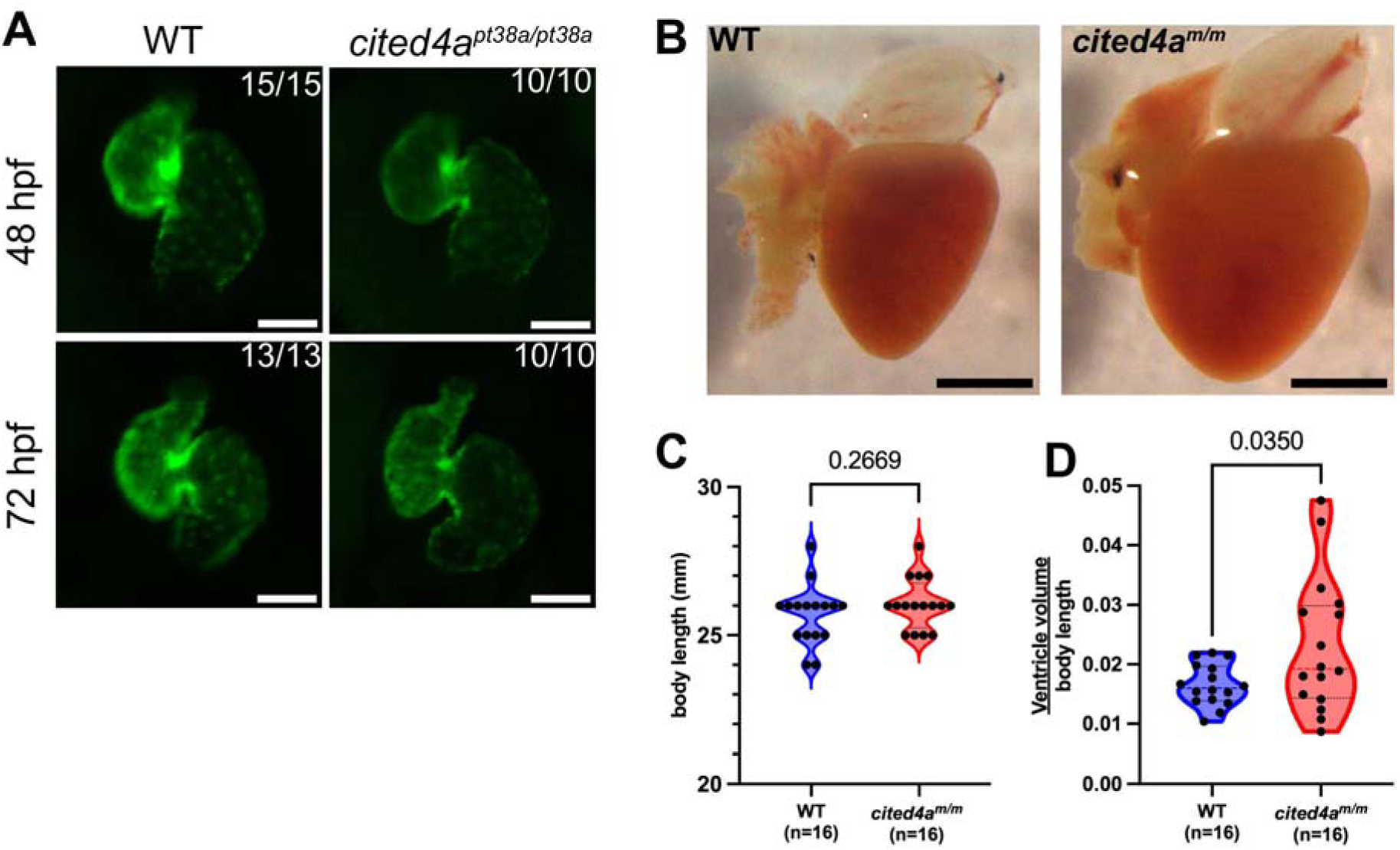
*cited4a* mutants heart development. (**A**) *Tg(myl7:EGFP);cited4a^pt38a/pt38a^* hearts at 48 and 72hpf. (**B**) *cited4a* mutant hearts are larger than WT hearts. (**C**) Quantification of WT and *cited4a* mutant body length and ventricular volume and statistical analysis using Unparied t test. White scale bar = 100µm. Black scale bar = 500µm.

**Supplementary Figure S6.**
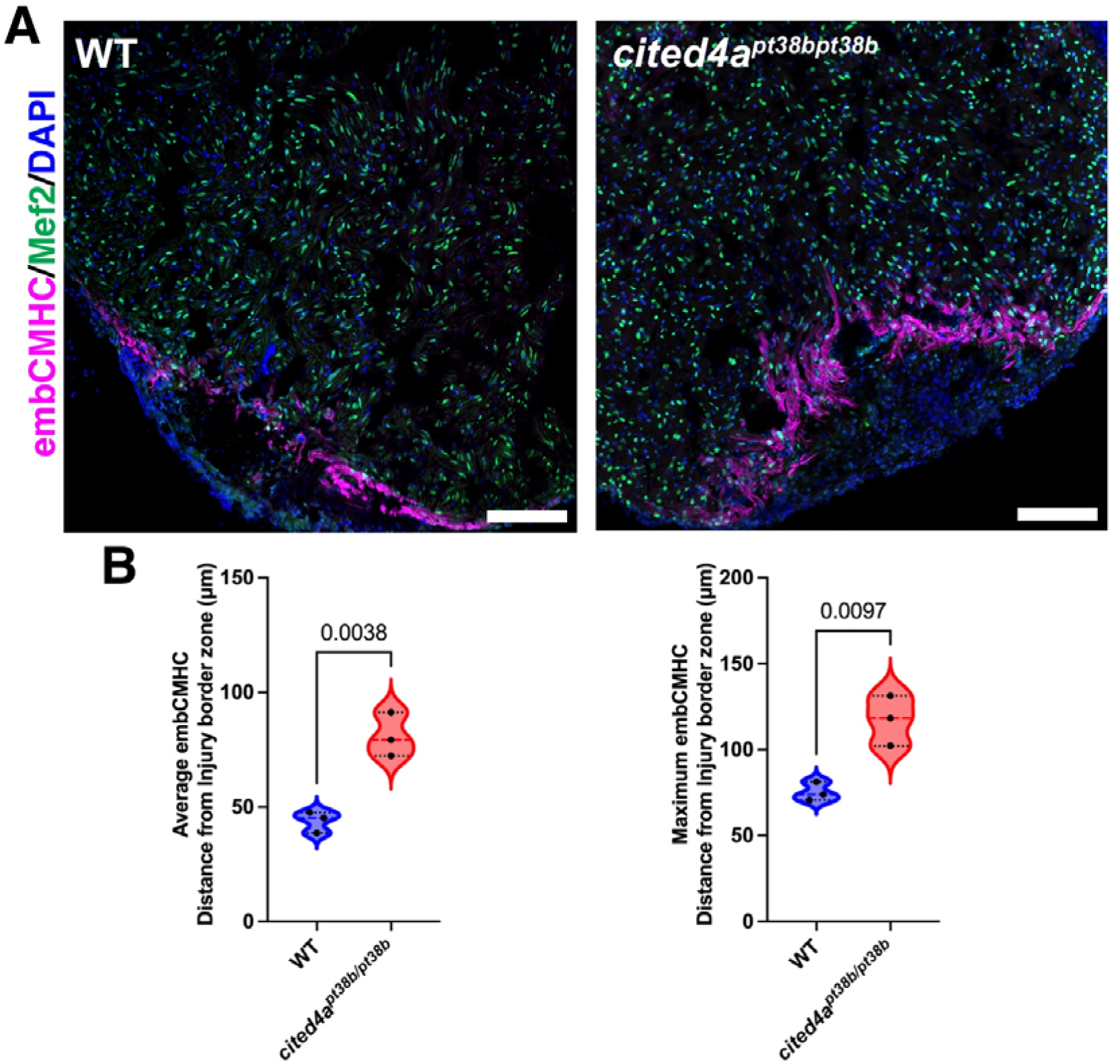
*cited4a^pt38b/pt38b^*mutant injured hearts show expansion of embCMHC expression in border zone CMs. (**A**) Representative images of WT and *cited4a^pt38b/pt38b^*, another mutant allele, injured 3dpa hearts stained with Mef2 and embCMHC. (**B**) Quantification of embCMHC expression in injured hearts using Unpaired t-test. Scale bar = 100µm.

**Supplemental Figure S7.**
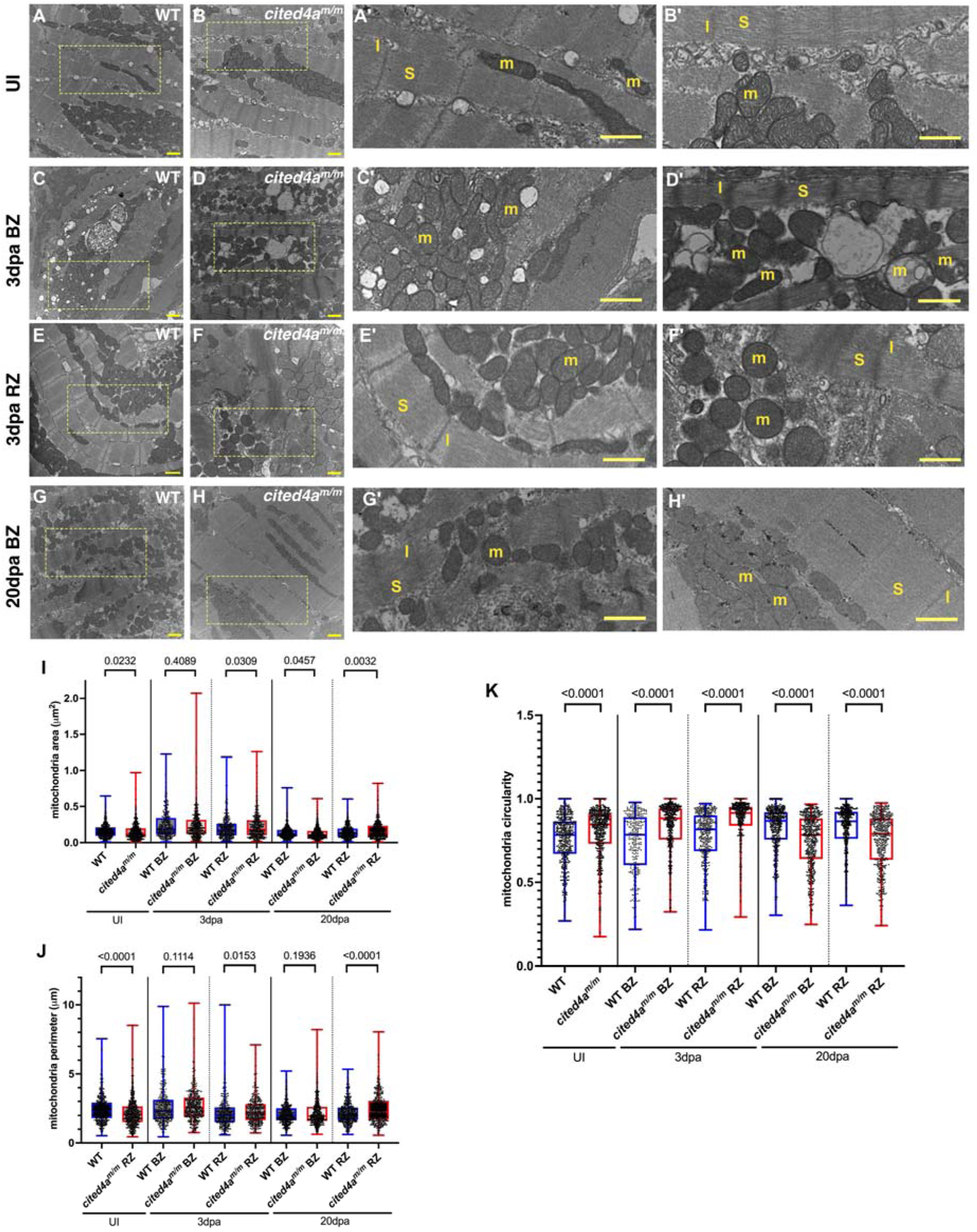
Transmission electron microscopy of WT and *cited4a* mutant hearts. Representative images of uninjured hearts show normal mitochondria and sarcomere structures in WT (**A, A’**) and *cited4a* mutant (**B, B’**) CMs. At 3dpa, *cited4a* mutant CMs show irregular sarcomere and mitochondria at the border and remote zones (**D, D’, F** and **F’**). At 20dpa, *cited4a* mutant CMs show normal sarcomere and mitochondria (**H, H’**) as compared to WT hearts (**G, G’**). Quantification of mitochondria area (**I**), perimeter (**J**) and circularity (**K**).

**Supplemental Figure S8.**
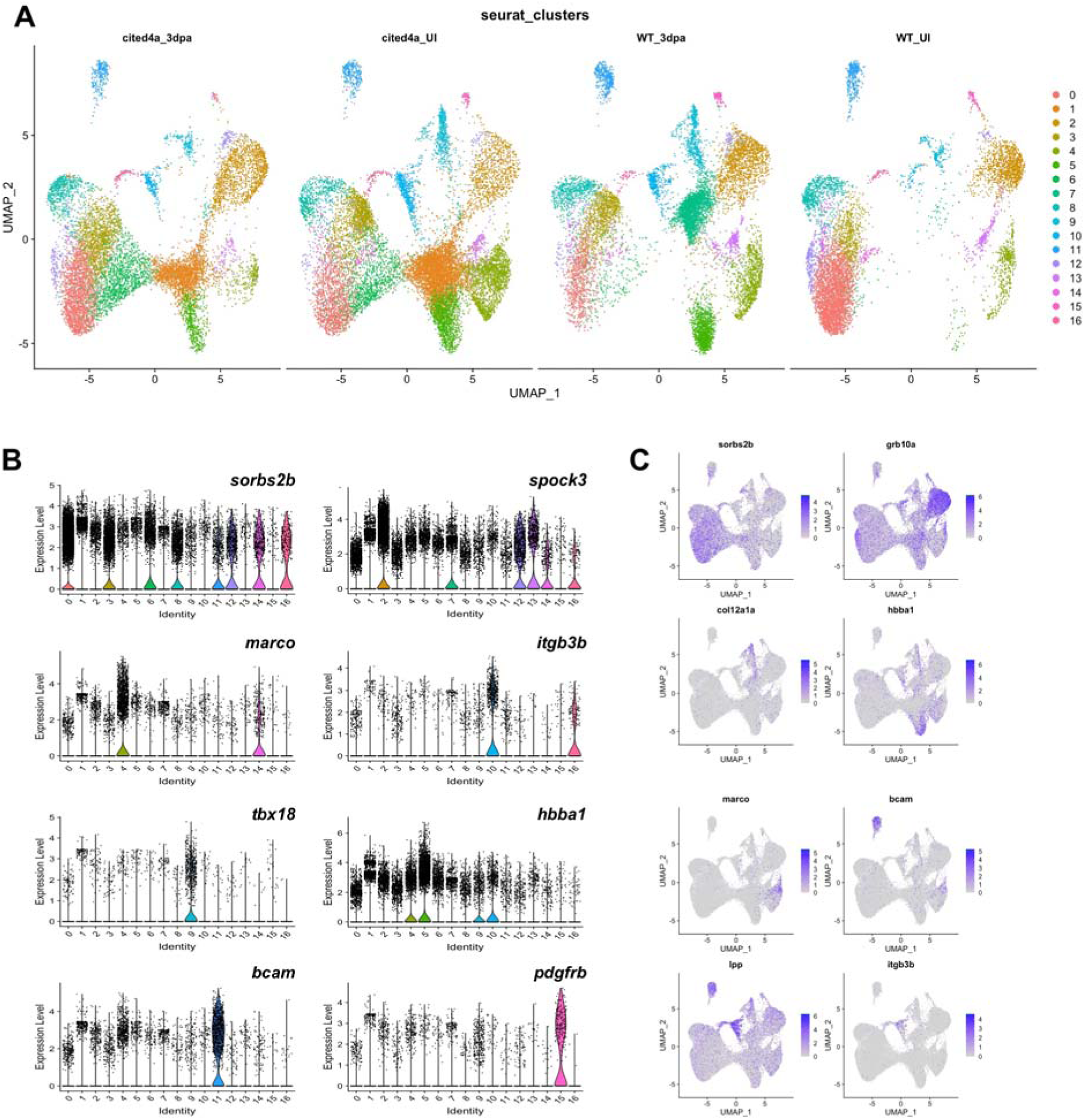
snRNA-seq analysis of *cited4a* mutant and WT hearts. (**A**) UMAP visualization of integrated *cited4a* mutant and WT hearts at 3dpa and uninjured split by condition. (**B**) Violin plots of genes from *cited4a* mutant and WT heart snRNA-seq. (**C**) Feature Plots of genes from this data set.

**Supplementary Figure S9.**
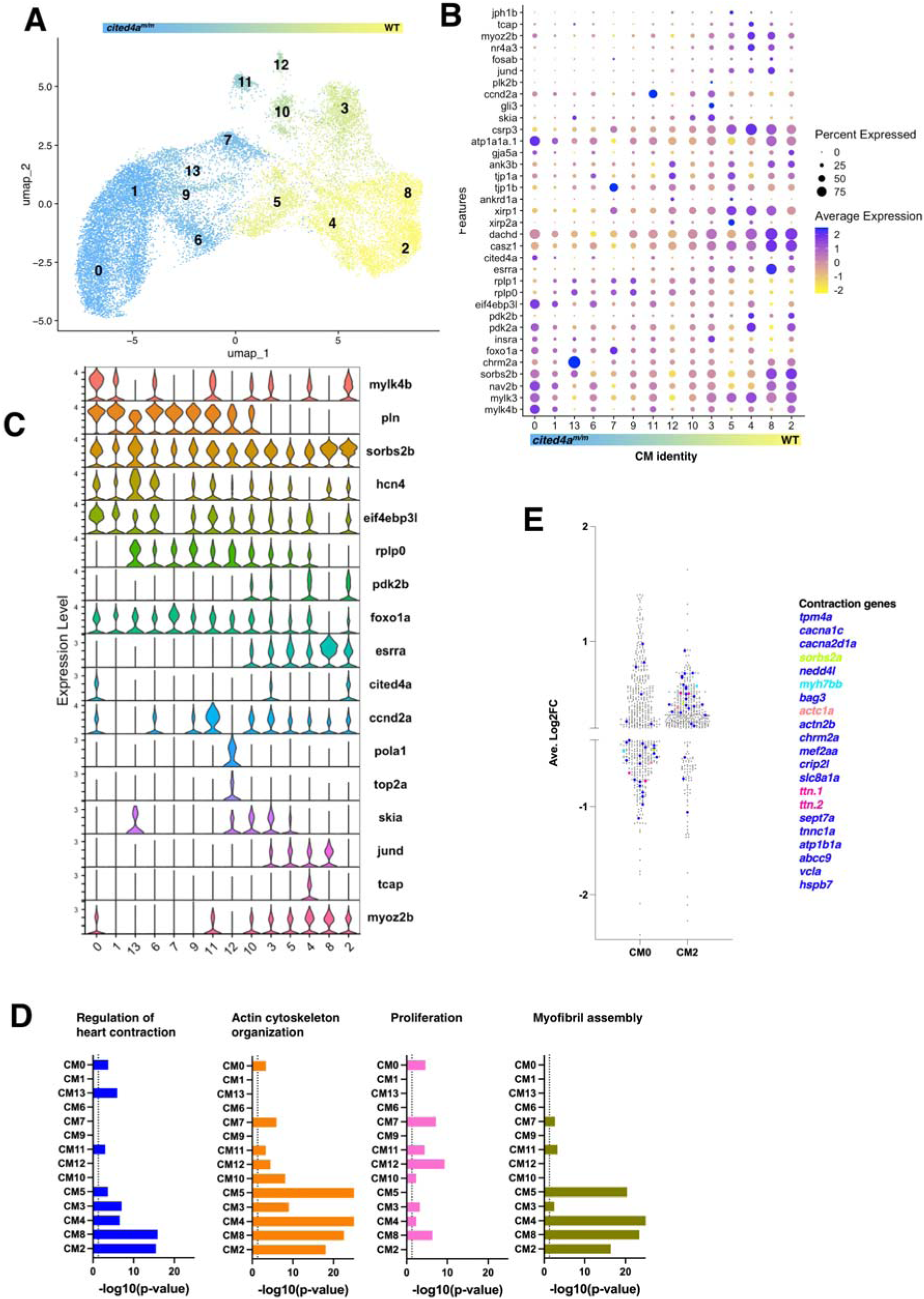
snRNA-seq analysis of CMs from *cited4a* and WT hearts. UMAP visualization of CMs color coded based on contribution of nuclei from WT and *cited4a* mutant hearts (**A**). Dot plot (**B**) and stacked violin plot (**C**) of CM genes. (**D**) Graphs showing GO biological process for each CM cluster. GO terms for regulation of heart contraction, Actin Cytoskeleton Organization, Proliferation and Myofibril Assembly are shown. (**E**) Comparison of cardiac contraction markers in CM0 and CM2. Representative genes are color coded to highlight the difference in their expression values between CM0 and CM2.

## Supplementary Tables

**Supplementary Table T1.** Marker genes from Seurat analysis of snRNA-seq for WT uninjured and 3dpa for clusters 0-14

**Supplementary Table T2.** Marker genes from Seurat analysis of reclustered CMs

**Supplementary Table T3.** TopFUN analysis of CM0-CM7 for GO:Biological Process

**Supplementary Table T4.** Bulk RNA-seq data of *cited4a* mutant and WT hearts at 3dpa with DAVID GO: Biological Process analysis of increased and decreased in *cited4a* mutant hearts.

**Supplementary Table T5**. Table showing Esrra target gene expression from the bulk RNA-seq data of decreased *cited4a* mutant hearts.

**Supplementary Table T6.** Marker genes from Seurat analysis of snRNA-seq for WT and cited4a mutant uninjured and 3dpa for clusters 0-16

**Supplementary Table T7.** Marker genes from Seurat analysis of snRNA-seq for WT and cited4a mutant uninjured and 3dpa for reclustered CM0-CM13

**Supplementary Table T8.** GO:Biological Process terms for positive markers from CM0-CM13 after harmony integration.

**Supplementary Table T9.** List of DNA primers, sgRNA, antibodies, and RNAscope probes used in this study.

