## Supplementary Table T5 for "Cited4a limits cardiomyocyte dedifferentiation and proliferation during zebrafish heart regeneration"

|  | **zebrafish gene/mammalian ortholog** | **Log2FC** | **FDR P-value** | **Evidence of regulation by ERR** |
| --- | --- | --- | --- | --- |
| sarcomere, cardiac myofibril assembly, cardiac muscle contraction | *actc1a*/*Actc1* | -2.857344 | 0.06268318 | [1-4] |
|  | *tnni1c/Tnni1* | -1.231056 | 0.08708434 | [2] |
|  | *atp1a1a.3/Atp1a1* | -1.231056 | 0.08708434 | [5] |
|  | *tcap/Tcap* | -1.053698 | 0.01062718 | [1, 5] |
|  | *casq2/Casq2* | -0.671384 | 0.047679 | [1, 2] |
| fatty acid metabolic process, fatty-acid beta-oxidation, fatty acid transport | *pm20d1.1/Pm20d1* | -6.636259 | 0.00080968 | [5] |
|  | *mlycd/Mlycd* | -1.789403 | 0.04859728 | [4] |
|  | *ucp3/Ucp3* | -1.156271 | 0.02019044 | [1, 2] |
|  | *fabp3/Fabp3* | -0.975798 | 0.00054025 | [1, 2, 5] |
|  | *acsl1b/Acsl1* | -0.922455 | 0.02583782 | [1, 2, 5] |
|  | *kcnj8/Kcnj8* | -0.920738 | 0.03760269 | [3, 5] |
|  | *cpt1b/Cpt1b* | -0.873331 | 0.00121145 | [1, 2, 5] |
|  | *acadvl/Acadvl* | -0.791339 | 0.05255644 | [1, 2] |
|  | *ech1/Ech1* | -0.760656 | 0.01882354 | [1, 2] |
|  | *acads/Acads* | -0.75527 | 0.00602896 | [4] |
|  | *acsl1a/Acsl1* | -0.74416 | 0.05478957 | [1, 2, 5] |
|  | *etfb/Etfb* | -0.713111 | 0.04007266 | [1, 2, 5] |
|  | *got2b/Got2* | -0.64747 | 0.06997834 | [5] |
|  | *hadhaa/Hadha* | -0.603006 | 0.04747693 | [1, 2] |
|  | *acaa2/Acaa2* | -0.588156 | 0.01203252 | [1, 2, 5] |
| mitochondria structure, function | *alas1/Alas1* | -1.670123 | 0.00099906 | [4] |
|  | *pdk2b/Pdk2* | -1.38142 | 0.06997834 | [5] |
|  | *ndufa4a/Ndufa4* | -1.150258 | 0.0031975 | [5] |
|  | *cox6b1/Cox6b1* | -1.0537565 | 0.00035139 | [1, 2, 5] |
|  | *cox4i1l/Cox4il* | -0.7532531 | 0.01819948 | [1, 5] |
|  | *zgc:110843/Cisd1* | -0.753038 | 0.03347193 | [4] |
|  | *mpc2/Mpc2* | -0.722044 | 0.04787357 | [1, 2] |
|  | *etfb/Etbf* | -0.713111 | 0.04007266 | [1, 2, 5] |
|  | *slc25a33/Slc25a33* | -0.66372 | 0.06804171 | [5] |
|  | *hccsb/Hccs* | -0.593581 | 0.04118405 | [5] |
|  | *atp5mc1/Atp5mc1* | -0.586759 | 0.01895478 | [1, 2] |
|  | *cox5b2/Cox5b* | -0.5854981 | 0.07601338 | [5] |
|  | *apooa/Apoo* | -0.582337 | 0.04854291 | [2, 5] |
| cardiac hypertrophy stress response | *tcap/Tcap* | -1.053698 | 0.01062718 | [1, 5] |
|  | *nppb/Nppb* | -0.935162 | 0.01460376 | [2] |
|  | *nppa/Nppa* | -0.690764 | 0.02188824 | [4] |
| transcriptional regulation | *esrra/Esrra* | -1.219838 | 0.00036952 | [1, 2, 5] |
|  | *jund/Jund* | -1.115503 | 0.07932112 | [4] |

Table S5

Representative shared decreased genes from *cited4a^pt38a^* 3 dpa bulk RNA-seq and ERR

deficiency models
